# Spatial profiling of the interplay between cell type- and vision-dependent transcriptomic programs in the visual cortex

**DOI:** 10.1101/2023.12.18.572244

**Authors:** Fangming Xie, Saumya Jain, Runzhe Xu, Salwan Butrus, Zhiqun Tan, Xiangmin Xu, Karthik Shekhar, S. Lawrence Zipursky

## Abstract

How early sensory experience during “critical periods” of postnatal life affects the organization of the mammalian neocortex at the resolution of neuronal cell types is poorly understood. We previously reported that the functional and molecular profiles of layer 2/3 (L2/3) cell types in the primary visual cortex (V1) are vision-dependent (Tan et al., *Neuron*, **108**(4), 2020; Cheng et al., *Cell*, **185**(2), 2022). Here, we characterize the spatial organization of L2/3 cell types with and without visual experience. Spatial transcriptomic profiling based on 500 genes recapitulates the zonation of L2/3 cell types along the pial-ventricular axis in V1. By applying multi-tasking theory (Adler et al., *Cell Systems*, **8**, 2019), we suggest that the spatial zonation of L2/3 cell types is linked to the continuous nature of their gene expression profiles, which can be represented as a 2D manifold bounded by three archetypal cell types (“A”, “B”, and “C”). By comparing normally reared and dark reared L2/3 cells, we show that visual deprivation-induced transcriptomic changes comprise two independent gene programs. The first, induced specifically in the visual cortex, includes immediate-early genes and genes associated with metabolic processes. It manifests as a change in cell state that is orthogonal to cell type-specific gene expression programs. By contrast, the second program impacts L2/3 cell type identity, regulating a subset of cell type-specific genes and shifting the distribution of cells within the L2/3 manifold, with a depression of the B-type and C-type and a gain of the A-type. Through an integrated analysis of spatial transcriptomic measurements with single-nucleus RNA-seq data from our previous study, we describe how vision patterns L2/3 cortical cell types during the postnatal critical period.

**Significance statement:** Layer 2/3 (L2/3) glutamatergic neurons are important sites of experience-dependent plasticity and learning in the mammalian cortex. Their properties vary continuously with cortical depth and depend upon experience. Here, by applying spatial transcriptomics and different computational approaches in the mouse primary visual cortex, we show that vision regulates orthogonal gene expression programs underlying cell states and cell types. Visual deprivation not only induces an activity-dependent cell state, but also alters the composition of L2/3 cell types, which are appropriately described as a transcriptomic continuum. Our results provide insights into how experience shapes transcriptomes that may, in turn, sculpt brain wiring, function, and behavior.

## Introduction

Early sensory experiences influence the development of neural circuitry throughout the mammalian brain (1–3). In the primary visual cortex (V1), visual experience is required to establish the circuitry for binocular vision (4, 5). Vision-dependent development of binocular circuits occurs in mice after eye-opening (postnatal day (P) 14). There are only a few binocular L2/3 neurons at eye opening and their number increases during the ensuing week. From P21 to P35, the tuning properties of these neurons improve markedly in a vision-dependent process. This developmental time window is referred to as the critical period (4, 6–8). Classically, the influence of early visual experience has not been examined at the level of the many cell types that form the building blocks of V1 circuitry. To address this gap, we previously performed single-nucleus RNA-seq (snRNA-seq) in normal- and dark-reared (NR, DR) mice during postnatal development, including the critical period (9). Using computational methods, we reconstructed the postnatal maturation of V1 cell types in NR mice and compared these profiles with DR mice. This enabled us to identify the cell types and gene expression programs impacted by vision. For most cell types, DR had little impact on molecular identity. However, DR had a pronounced effect on glutamatergic cell types in the supragranular layers (L) 2/3/4. In NR mice, L2/3 glutamatergic neurons comprise three molecularly distinct types (A, B, and C). These L2/3 types occupy three partially overlapping sublayers along the pial-ventricular axis of L2/3 (upper, middle, and lower, respectively) based on the *in situ* expression patterns of three marker genes. DR altered the expression patterns of these marker genes, and a comparison of transcriptome-based clusters in DR did not correspond to NR types. Based on these observations, we concluded that visual deprivation during the critical period selectively disrupted L2/3 cell types. Furthermore, visual deprivation regulated large groups of genes, including determinants of cell adhesion and synapse formation. However, as clusters in DR could not be matched to NR types, we could not separate vision-dependent changes in cell state from vision-dependent changes in cell type, their organization, or both.

Here, we combined spatial transcriptomics and computational analyses to map the organization of L2/3 cell types in V1, and to assess the impact of visual deprivation on this organization. The gene expression changes due to visual deprivation comprise two independent transcriptomic programs. The first program is upregulated across multiple cell types throughout the visual areas and represents an activity-dependent cell state. The second program regulates a subset of cell type-specific genes within L2/3, and its impact on cell type composition can be interpreted using multi-tasking theory (10), a recently proposed framework for analyzing gene expression continua. Thus, visual deprivation affects both activity-dependent transcriptomic states and cell-type composition in L2/3.

## Results

### Spatial transcriptomic analysis of V1 reveals the continuous sublayered arrangement of L2/3 glutamatergic cell types

We first sought to explore the spatial arrangement of L2/3 cell types. To do this, we performed multiplexed error-robust fluorescence *in situ* hybridization (MERFISH (15)) on brain coronal sections containing V1 obtained from normally-reared (NR) mice at postnatal day (P) 28 (**Figure 1A**). MERFISH is an imaging-based approach that allows simultaneous mapping of hundreds of genes in individual cells within an intact tissue section. Using computational methods, cells can be classified into types based on combinatorial gene expression patterns, and the spatial arrangement of each cell type can be visualized. For these experiments, we selected a panel of 500 genes that included cortical area- and layer-specific markers, marker genes for all major cortical cell populations (9, 13), and activity-regulated genes (11) (**Tables S1, S2**). Notably, given our specific goal of studying cell type-specific changes within L2/3, we included 170 genes selected from the 286 L2/3 cell-type-identity genes we previously reported (9) (see **Table S1**).

**Figure 1.**
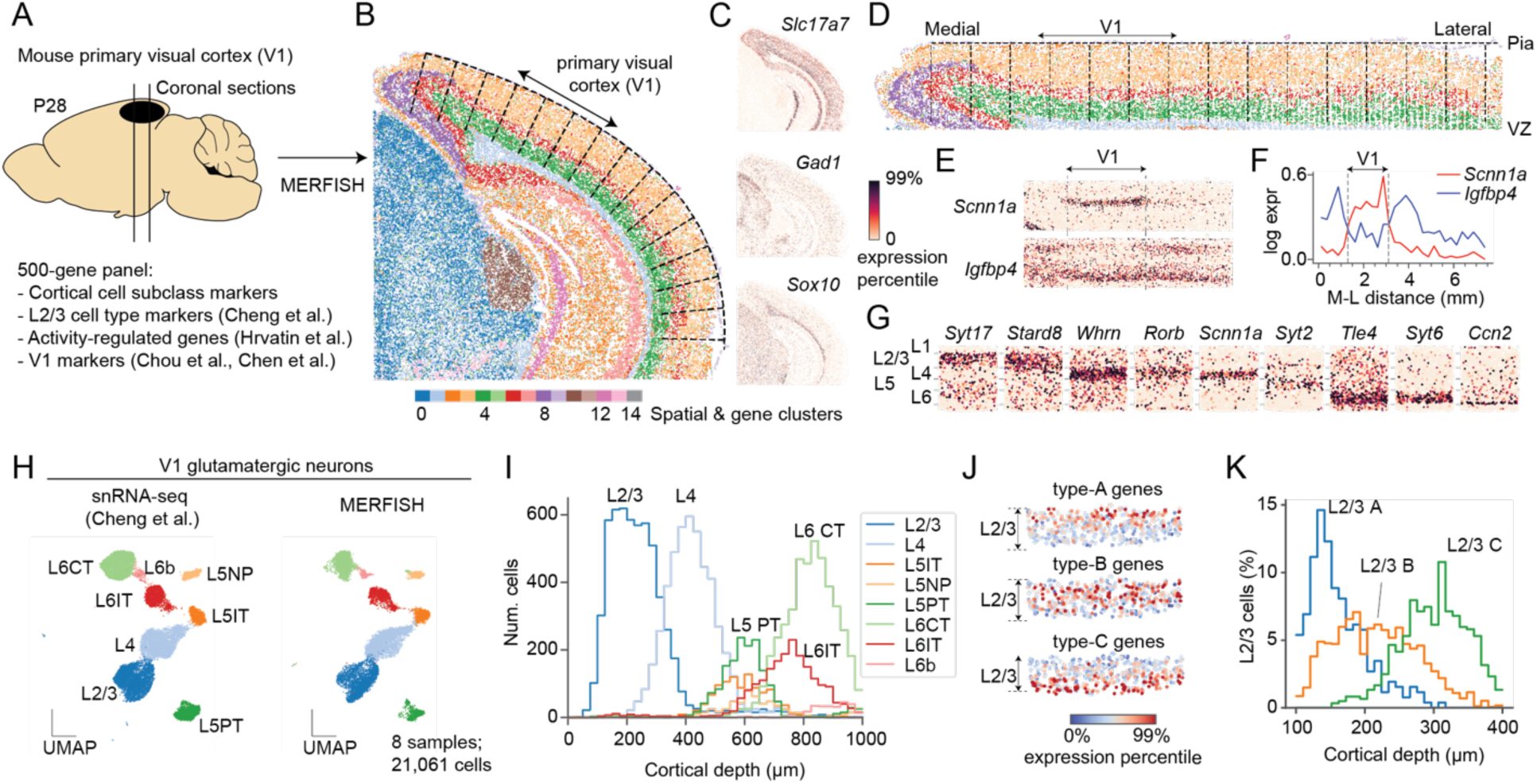
MERFISH recapitulates the spatial organization of L2/3 glutamatergic neurons in V1. **(A)** MERFISH was performed on mouse coronal brain sections using a panel of 500 genes. The panel was designed to resolve cell types and cell states in V1 based on published studies (9, 11–13). **(B)** An overview of the MERFISH data, showing a thin (10 µm) coronal section including the neocortex, parts of the hippocampus, and midbrain structures. Individual cells are colored by their cluster membership based on both gene expression similarity and spatial proximity (see **Figure S1A**). **(C)** *In situ* gene expression patterns of *Slc17a7,* a glutamatergic neuron marker; *Gad1,* a GABAergic neuron marker; and *Sox10*, an oligodendrocyte marker. Colors represent gene expression levels going from 0 to 99 percentile. **(D)** Cortical cells with defined locations along the medial-lateral (M-L) and pial-ventricular (P-V) axis of the cortex. Tissue was straightened *in silico* to facilitate downstream analysis. Cells are colored as in panel B. **(E)** Two genes, *Scnn1a* and *Igfbp4* with area-specific *in situ* signatures. **(F)** Expression levels of genes in panel E along the M-L axis. The location of V1 is highlighted. **(G)** Examples of genes with layer-specific signatures in V1. **(H)** Subclasses of V1 glutamatergic neurons are represented in UMAP embeddings obtained from integrating snRNA-seq (left panel; (9)) and MERFISH (right panel; this study) using Harmony (14). **(I)** Spatial distributions of glutamatergic subclasses along the cortical depth (P-V) axis. **(J)** Aggregated expression in V1 L2/3 for gene groups defining types A (n=64 genes), B (n=35 genes) and C (n=71 genes). Type A, B, and C genes are distributed in upper, middle, and lower L2/3, respectively. Colors represent gene expression levels going from 0 to 99 percentile. **(K)** Spatial distributions of types A, B, and C along the cortical depth spanning L2/3. MERFISH cell labels were transferred from previously published snRNA-seq data (9).

We performed a preliminary clustering of the cells from the entire section, defining clusters using a combination of molecular similarity and spatial proximity (**Methods**). This analysis identified 15 cell clusters, which highlighted different high-level anatomical structures of the mouse brain, including the neocortex with its characteristic laminar structure, parts of the hippocampus, and midbrain structures (**Figures 1B, S1A**). These associations were supported by *in situ* expression patterns of well-known genes. For example, *Slc17a7* (*Vglut1*), a marker for glutamatergic neurons, is prominently expressed in the cortex and parts of the hippocampus, two regions where glutamatergic neurons comprise 80∼90% of all neurons (16–18) (**Figure 1C**). In contrast, *Gad1*, a marker for GABAergic neurons, is expressed at low levels in the cortex but high levels in the midbrain, where inhibitory neurons represent ∼50% of neuronal cells (17, 18). *Sox10*, a marker for oligodendrocytes, is enriched in the white matter beneath the cortex.

We used the pial surface to define the tangential (medial-lateral) and vertical (pial-ventricular) coordinates of each cell and demarcated areas of the neocortex (**Figures 1B-G, S1B**). We localized the primary visual cortex (V1) based on the enrichment of *Scnn1a* and the depletion of *Igfbp4* along the tangential axis. Both markers, described in recent reports (12, 13), allowed us to identify V1 as a ∼2mm wide x 1mm deep region (**Figures 1E,F**).

We used an integrative approach to classify each V1 cell in MERFISH into one of three classes (excitatory neurons, inhibitory neurons, and non-neuronal cells) and each neuron into one of 12 subclasses, as in Cheng et al. (9) (**Figures 1H, S1C-E**). The relative frequencies of the subclasses tightly corresponded between MERFISH and snRNA-seq (Spearman correlation = 0.95), and the different subclasses of glutamatergic neurons were spatially localized within expected layers along the pial-ventricular axis (**Figures 1I, S1C-F**).

Previously, we found that L2/3 cells can be clustered into three types (L2/3A, L2/3B, and L2/3C). Individual marker genes for these types – *Cdh13* for L2/3A, *Trpc6* for L2/3B, and *Chrm2* for L2/3C - were expressed in upper, middle, and lower regions of L2/3 along the pial-ventricular axis (9). However, the MERFISH data allowed us to validate the three-layered zonation of L2/3 types using a much larger gene panel (**Figures 1J, S2**). Combinatorial gene signatures based on 170 type-identity genes enabled classification of MERFISH L2/3 cells as L2/3A, L2/3B, or L2/3C. As expected, soma of the three cell types localized to the upper, middle, and lower sublayers within L2/3 (**Figure 1K**). Type A and C identity genes were expressed in a more restricted spatial pattern, whereas type B identity genes were expressed more broadly.

### Multi-tasking theory relates the spatial zonation of L2/3 glutamatergic neuronal types to their continuous transcriptomic variation

The gene expression patterns that underlie the transcriptomic continuum are most evident when we analyzed the data using diffusion pseudotime (DPT) (19), an approach based on diffusion maps (20). For snRNA-seq, DPT orders cells according to their type identity along the x-axis and their corresponding type-identity genes along the y-axis (**Figure 2A**). The genes are expressed at different levels in a graded fashion, with no clear boundaries between the domains of each cell type. DPT produces a similar result for the MERFISH dataset, with cells now ordered based on their location along the cortical depth **Figure 2B**), further supporting the link between continuous transcriptomic variation and spatial zonation.

**Figure 2.**
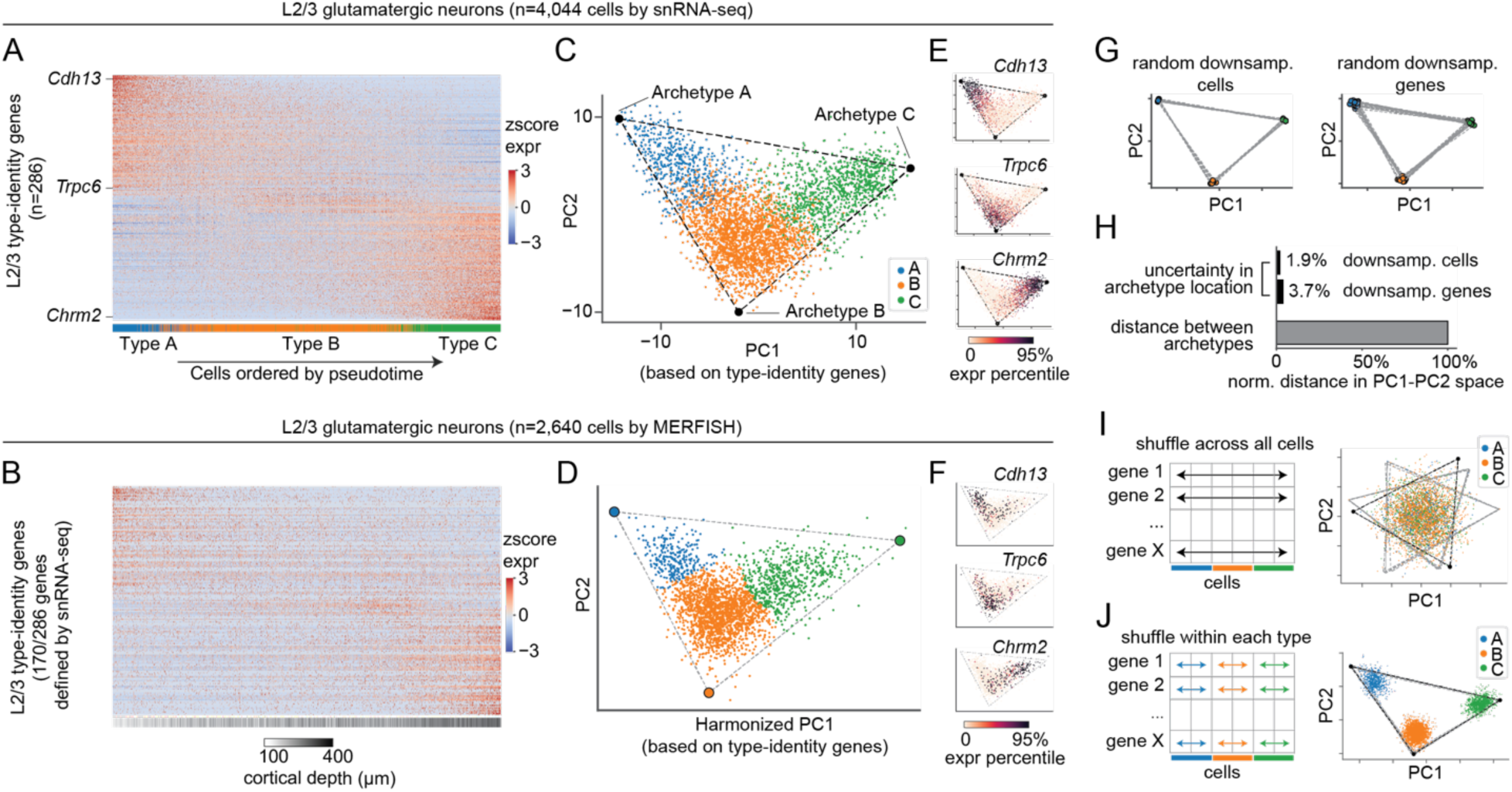
L2/3 glutamatergic neurons form a continuous manifold bounded by a triangle, whose vertices represent archetypes. **(A-B)** Expression heatmaps of type-identity genes (**Table S1**) across L2/3 glutamatergic neurons for snRNA-seq (A) and MERFISH (B). Expression was quantified as z-scored, log- and size-normalized counts. Cells (x-axis) were ranked by diffusion pseudotime (DPT) and colored by cell-type identity in (A) and cortical depth in (B) (annotation bar, bottom). **(C-D)** L2/3 cells represented in PC1 and PC2 space based on the same type-identity genes in (A-B). Cells are colored by type. The bounding triangle was inferred using computational procedures described in Adler et al. (10) (see Methods). **(E-F)** Expression levels of *Cdh13*, *Trpc6,* and *Chrm2* in PC1/PC2 space for scRNA-seq (E) and MERFISH (F). **(G-H)** Stability of the bounding triangle over randomized trials involving random down-sampling of the original dataset to 80% of cells (G, left panel) or 80% of genes (G, right panel). Comparison % variation in archetype coordinates during randomizations relative to the typical inter-archetype distances in these trials. Distances are measured by Euclidean distance in PC1-PC2 space. (H). **(I-J)** L2/3 cells in PC1/PC2 space after shuffling each gene independently across all cells (I), and after shuffling each gene independently across cells within each type (J), respectively. The gray lines represent triangular fits using 80% of cells, randomly down-sampled ten times independently. Thus, these cells form a continuum of types rather than three noisy cell types (see Figure S4). Panels G-J used snRNA-seq data only.

Strikingly, such a link is predicted by a recently proposed framework known as multi-tasking theory (10). This theory posits that *“if gene expression lies on a D-dimensional manifold (e.g., D=3 for a tetrahedron, D=2 for a triangle, or D=1 for a line), the cells should show at least a D-dimensional spatial zonation [in tissue]”* (10). Applying this framework to our data, we find that L2/3 neurons lie on a V-shaped manifold along the first two principal components (PCs), which is bounded by a triangle (**Figure 2C; Methods**). The structure of the manifold and the coordinates of the bounding triangle are similar for both the snRNA-seq and the MERFISH datasets (**Figures 2D, S3**). According to multi-tasking theory, vertices of the triangle represent “archetypal” cell types, with the gene expression profile of each archetype specialized for a particular function (see below). Indeed, the three vertices map to the L2/3 A, B and C types identified by clustering, and cells inside the triangle are mixtures of the archetypal gene expression profiles (**Figure 2E,F**). Finally, consistent with the theory, the organization along the manifold from A to B to C is mirrored in the spatial zonation along the pial-ventricular axis (**Figure 1J,K**).

We performed additional analyses to assess the robustness of the V-shaped manifold and the bounding triangle. Repeating the analyses with subsets of cells or genes resulted in only minor perturbations of the bounding triangle (**Figure 2G,H**). Repeating the analyses with shuffled versions of the original gene expression matrix further substantiated the continuous nature of the transcriptomic variation. First, we shuffled gene expression values independently for each gene across L2/3 cells, which preserves individual gene expression distributions while disrupting their correlations. This resulted in all cells collapsing towards the center and a total disruption of the triangular structure (p<0.001; t-ratio test) (**Figure 2I, S4**). We then shuffled gene expression values independently for each gene within each cell type (**Figure 2J**). This procedure distinguishes between two scenarios: a genuine continuum vs. discrete types seemingly continuous due to noise in the data, which could arise from the intrinsic stochasticity of gene expression or sampling noise in single-cell sequencing (21). When applied to clusters that span a continuum, the shuffling procedure splits continua into discrete clusters as long as the level of noise in the data are low to intermediate (**Figures S5A,C,E**). In contrast, when the clusters are already discrete it has no effect regardless of noise level (**Figures S5B,D,F**). Applying this shuffling procedure to our data splits L2/3 cells into three clusters (**Figure 2J**), supporting that transcriptomic continuum is genuine and not an artifact of noise.

What is the functional significance of this continuum? According to multi-tasking theory, such transcriptomic continua arise as an optimal solution for division of labor among cells in tissues where the a cell’s functional identity is related to its spatial position. While the theory does not specify these functions, examining the genes that define the archetypes may provide some clues. By performing gene ontology (GO) analyses, we found that archetype-defining genes are enriched for biological processes related to neuronal development and functions (**Figure S6**).

Programs involved in neural wiring, including “axon guidance” and “cell-cell adhesion,” show up in all three types (A, B, and C). For example, among known cell-recognition molecules related to neural wiring, type A expresses *Cdh13, Cntn5, Epha6, Sema6a* and *Robo1*; type B expresses *Epha3* and *Sema4a;* and type C expresses *Cdh12, Cntn2* and *Robo3* (**Table S3**). The differential expression of these genes may be related to the distinct projection patterns of A, B and C types to the higher visual areas (9, 22). Other programs, such as “sensory perception of light stimulus” and “detection of mechanical stimulus involved in sensory perception,” are specific to types B and C, respectively. Although one can only speculate on the functional meaning at this stage (see **Discussion**), the manifold representation of L2/3 neurons proves useful in understanding global and cell type-specific changes due to visual deprivation, which we analyze in the next section.

### Transcriptomic heterogeneity in vision-deprived L2/3 neurons involves two distinct gene programs

To characterize the changes induced by visual deprivation, we obtained MERFISH data from mice dark-reared (DR) between P21-P28, following our published experimental protocol (**Figure 3A; Methods**) (9). Comparisons of DR and NR sections collected at P28 showed an upregulation of 13 genes (FC>2, FDR < 0.05, **Figure S7A**), a small subset of vision-dependent genes detected by snRNA-seq (see below). All 13 genes were canonical immediate early genes (IEGs) (11), and their upregulation was localized to the visual areas (**Figure 3B**). Along the tangential axis, IEG levels were significantly higher in V1 than in its flanking regions (**Figures 3C,D, S7B**). Consistent with prior snRNA-seq data, we observed that the up-regulation of IEGs was broadly shared by all neuronal subsets and, therefore, visible throughout the cortical depth (**Figures 3C,E, S7C**). Within L2/3, canonical IEGs, such as *Fos*, *Nr4a2, Junb and Egr4*, were up-regulated in DR similarly in all sublayers (**Figures 3F, S8A,C**).

**Figure 3.**
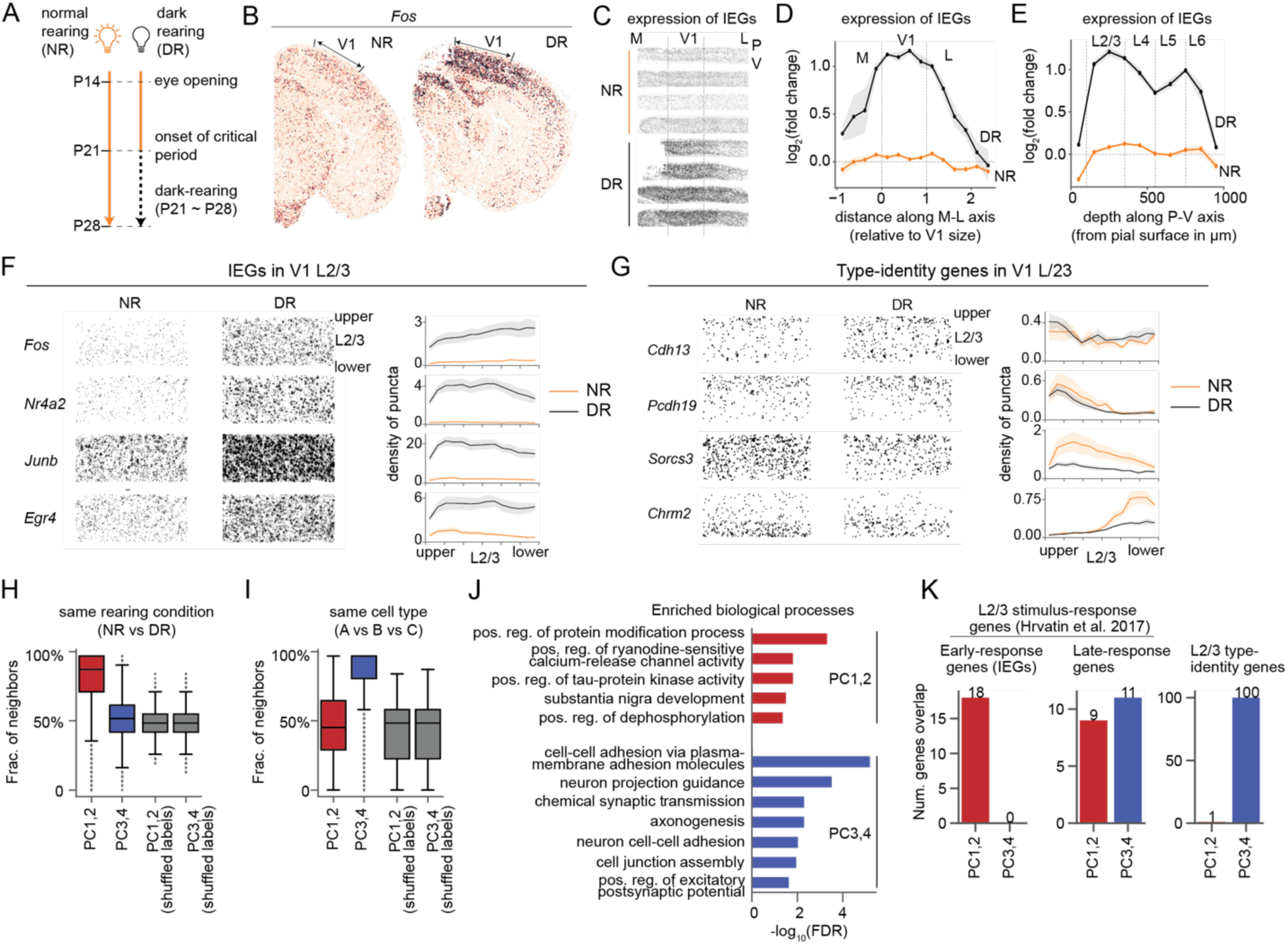
Gene programs driving transcriptomic heterogeneity in normal- and dark-reared L2/3 neurons. **(A)** Protocol for dark-rearing (9). **(B)** *In situ* expression patterns of *Fos* in NR and DR coronal sections. Locations of V1 are highlighted. **(C)** Aggregated *in situ* expression of IEGs (11) in NR and DR in V1 and its flanking cortical regions. Rows in each condition correspond to biological replicates. **(D-E)** Aggregated expression of IEGs along the M-L axis (D) and the P-V axis (E). Fold change is quantified relative to the mean expression level over the entire V1 region for all NR samples. **(F,G)** *In situ* expression of representative IEGs (F) and type-identity genes (G). Individual dots are transcript molecules detected by MERFISH. Line plots show the distribution of puncta along the P-V axis from upper to lower L2/3. The density of puncta were defined as the number of puncta per 100 µm^2^ area and were averaged across samples. Error bars represent standard error of the mean. **(H-I)** Distribution of L2/3 cells’ neighbor identities in terms of the the fraction of k nearest neighbors (k=30) that are of the same rearing condition (NR vs DR; panel H) and of the same type (A vs B vs C; panel I). PCs are computed from n=6,360 HVGs (see Methods and Figure S9). **(J)** Gene ontology (GO) analysis shows biological processes enriched in genes driving PC1-PC2 (in red) and PC3-PC4 (in blue). Raw results from the enrichment analysis were filtered to remove redundant terms (see Methods). A full list of the top 10 significant GO terms with FDR < 0.05 is shown in **Figure S9E**. Top 100 genes for each PC ranked by the absolute value of PC loadings are used. **(K)** Overlap between PC-driving genes and previously defined gene groups. Activity-regulated genes, including early response genes (immediate early genes - IEGs) and late-response genes in L2/3 glutamatergic neurons are from Ref (11). Type-identity genes from Ref (9) are listed in Table S1.

We previously reported that DR selectively disrupts the transcriptomic signatures of L2/3 cell types in single-nucleus (sn) RNA-seq data from V1 (9). Our results were based on unsupervised clustering of snRNA-seq profiles from NR and DR mice. Most cell types correspond 1:1 between NR and DR with the notable exception of L2/3 neurons. In the MERFISH data, while type-identity genes by and large maintained their sublayered expression, they also exhibited some vision-dependent alterations in DR. L2/3A-specific genes were relatively unchanged, but L2/3B- and L2/3C-specific genes were downregulated (**Figures 3G, S8B-D**). Taken together, **Figures 3A-G** suggest that vision impacts L2/3 transcriptomes in two distinct modes, one comprising broadly expressed genes (e.g. IEGs), and the other impacting genes associated with cell type identity (**Figure S8E,F**).

These results motivated a re-analysis of our published snRNA-seq dataset (9). We combined NR and DR datasets at P28 and applied principal component analysis (PCA) using 6,360 highly variable genes (HVGs; see Methods). We focused on the top four PCs, which exhibited a clear spectral gap from other components (**Figure S9A**). PC1 and PC2 separated cells by rearing condition (NR vs. DR mice), while NR and DR cells intermixed in the PC3-PC4 space (**Figures 3H, S9B-C**). Conversely, PC3-PC4, but not PC1-PC2, separated NR cells by type identity (**Figures 3I, S9B-C**), recapitulating the triangular manifold in **Figure 2B**, which was estimated using 286 type-identity genes. Thus, transcriptomic variation in L2/3 glutamatergic neurons is driven by two gene programs: the first, encoded in PC1-PC2, captures vision-dependent changes, while the second, encoded in PC3-PC4, captures cell type identity.

As PCs are derived from orthogonal directions in transcriptomic space, we hypothesized that the gene programs encoded in PC1-PC2 vs. PC3-PC4 represent distinct biological processes. Gene ontology analysis shows that genes driving PC1 and PC2 were enriched in the regulation of protein modification and signaling pathways (**Figures 3J, S9E**), likely reflecting shifts in cell state due to experience-dependent activity. By contrast, PC3 and PC4 were enriched for genes associated with cell-cell adhesion, axonogenesis, neuron projection guidance, and chemical synaptic transmission (**Figures 3J, S9E, Table S4**), which are associated with neuronal cell-type identity (23–26). We also compared the top-loading genes within each PC with a list of known activity-regulated genes in L2/3 glutamatergic neurons from Hrvatin et al. (11). This list includes 42 “early-response” genes, among which many are canonical IEGs, that were conserved across cell types in the original study, and 37 “late-response” genes, which were largely cell type-specific (**Table S5**). PC1-PC2 genes strongly overlapped with IEGs (odds ratio = 25, p<10^-10^; Fisher’s exact test; significant enrichment) and barely overlapped with type-identity genes (odds ratio = 0.1, p=0.002; Fisher’s exact test; significant depletion) (**Figures 3K**). By contrast, PC3-PC4 genes did not contain IEGs and significantly overlapped with type-identity genes (odds ratio = 40, p<10^-10^). Late-response genes featured in equal numbers across PC1-PC2 and PC3-PC4.

These results highlight the consistency between the MERFISH and the snRNA-seq datasets and present the following overall picture. The transcriptomic variance in NR and DR layer 2/3 cells can be decomposed into two independent programs: (1) PC1-PC2, associated with experience-regulated cell states, and (2) PC3-PC4, associated with cell type identity. However, we find that the contribution of these programs to the overall variance within a dataset differs by rearing condition. For NR, the projected variance along PC3-PC4 is 2.9-fold higher than that along PC1-PC2. By contrast, for DR, the projected variance along PC1-PC2 is 2.8-fold higher than along PC3-PC4. This disparity, however, drives the lack of correspondence in the HVG-based clustering of NR and DR L2/3 neurons reported in Cheng et al (9). Consistent with this, a “focused” clustering of DR L2/3 neurons using the 286 type-identity genes recovers cell clusters corresponding 1:1 with types A, B and C in NR (**Figure S10**). Overall, this suggests that despite being masked by experience-dependent changes, cell identity programs persist in DR (**Figure S9D**), albeit with major alterations (**Figure 3G**). We now analyze these alterations.

### Visual deprivation alters the composition of L2/3 cell types along the transcriptomic continuum

We used the 2D manifold representation (**Figure 2C,D**) to study cell type-specific changes induced by visual deprivation. We hypothesized that vision could regulate one or more of the following features: a) the locations of the archetypes, i.e., the bounding triangle; b) the distribution of cells along the manifold, which correspond to changes in cell type composition and spatial zonation; and c) the expression levels of one or more type-identity genes.

To evaluate whether archetype coordinates change, we recalculated PCs based on the 286 type-identity genes as in **Figure 2C**, but now using both NR and DR cells from the published snRNA-seq data. Using this representation, we separately inferred archetype coordinates using cells from each biological sample (n=4 replicates for each condition, including P38 data from ref (9)). These calculations verified that L2/3 cells from NR and DR mice occupy similar triangular regions, with no significant changes detected in archetype coordinates (3∼10% changes relative to the minimum distance between archetypes) (**Figure S11A,C**).

Individual cells, however, distributed differently between NR and DR, shifting away from the region proximal to archetype B towards archetype A, and away from archetype C in DR (**Figure 4A; Figure S11B**). This reorganization between NR and DR was also predicted by two alternative methods: optimal transport analysis (27, 28) (**Figure 4B; Methods**) and supervised classification-based label transfer from NR to DR cells (**Figure 4C**). The compositional difference between NR and DR at P28 or P38 was larger than the difference between P28NR and P38NR, or between P28DR and P38DR (**Figure S11D**). This suggests that the effect of DR on L2/3 transcriptomic identities is consistent over at least ten days. Furthermore, similar changes in cell type composition were also recovered in the MERFISH data (**Figure S12**). Additionally, MERFISH shows that the overall expression levels of type B and C genes decrease in DR, especially in middle and lower L2/3 (**Figure 4D,E; Figure S13**). By contrast, the expression of type A genes is relatively unchanged and consistently enriched in upper L2/3.

**Figure 4.**
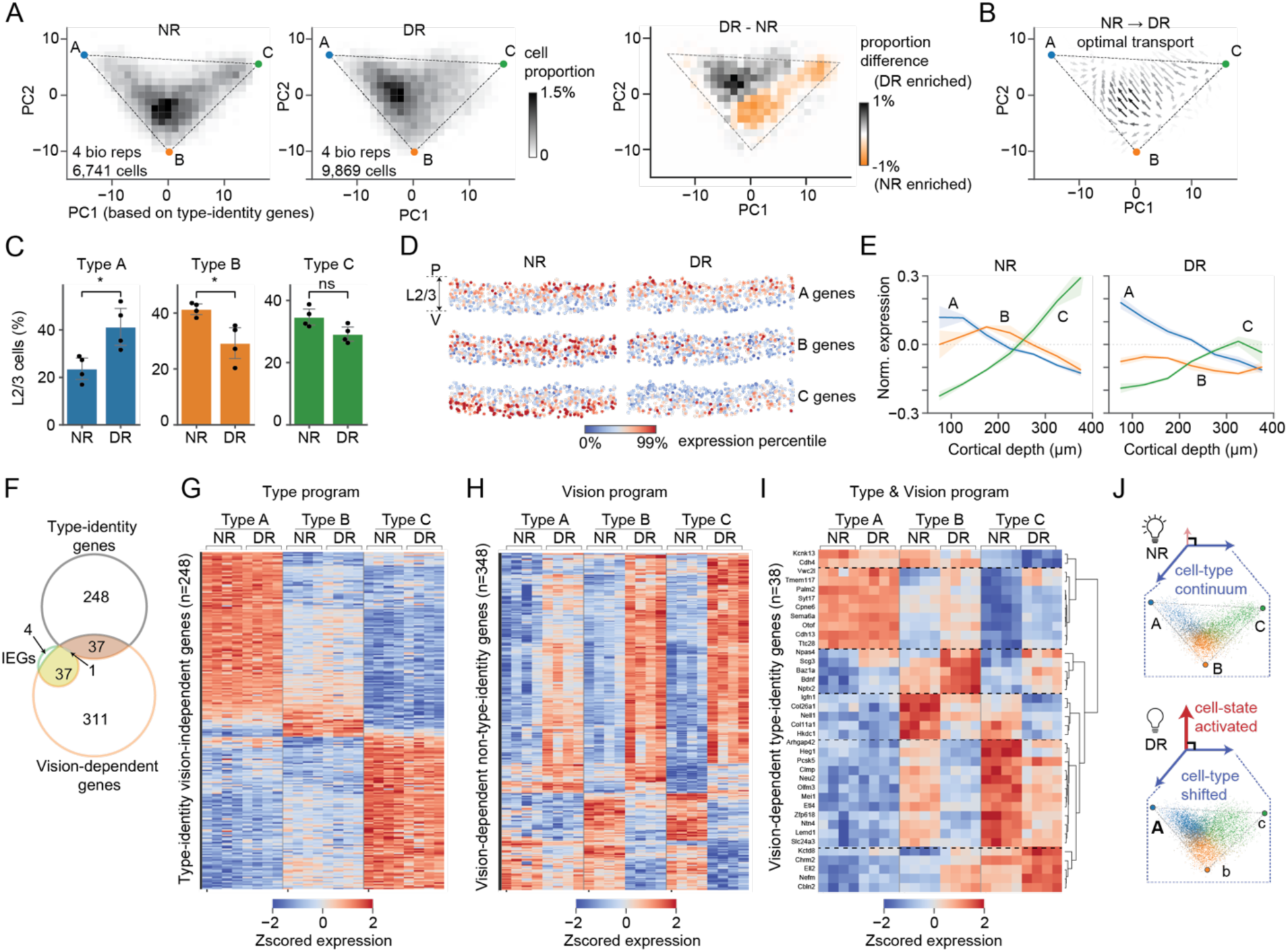
Visual deprivation redistributes cells within the L2/3 type triangle. (A) Distribution of L2/3 types within the L2/3 type-triangle. Cell density for NR cells (left), DR cells (middle), and the difference between NR and DR (right) are shown in PC1 and PC2 space derived from type-identity genes. (B) Optimal transport analysis (27, 28) showing the transport map connecting the NR to DR distribution. The arrows indicate the direction of local redistribution of cells in the reduced gene expression space. (C) The proportions of L2/3 cells assigned as types A, B and C for NR and DR samples. The type assignment is based on type A, B and C identity gene expression (see Methods). (D) *In situ* expression levels of A, B and C type-identity genes for NR and DR samples. (E) Expression of A, B and C identity genes along the P-V axis from upper to lower L2/3 for NR (left panel) and DR (right panel) samples. Mean expression levels were shown relative to the baseline levels, which are defined as the mean expression levels in NR samples averaged across L2/3. (F) Overlap between vision-dependent genes, type-identity genes and previously identified L2/3 IEGs (11). (G-I) Expression profiles of type-identity vision-*in*dependent genes (G), vision-dependent *non*-type-identity genes (H), and vision-dependent type-identity genes (I). Expression levels are z-score normalized across all samples independently for each gene. Gene numbers correspond to panel (F). (J) Diagram shows L2/3 transcriptomes comprise two orthogonal axes of variations – cell-state programs and cell-type programs. Visual deprivation activates cell-state programs and shifts cell-type programs away from archetypes B and C towards A.

We performed differential gene expression analysis between NR and DR samples to identify vision-dependent genes within each type. In total, we identified 386 unique vision-dependent genes, of which 70 were regulated in type A, 226 in type B, and 304 in type C (fold change >2 and FDR <0.05). These genes included 90% (n=38/42) of previously identified L2/3 IEGs (11), while the IEGs represent ∼10% (n=38/386) of the vision-dependent genes (**Figure 4F**). Moreover, most vision-dependent genes (90%; n=348/386) are not type-specific, and they were up- or down-regulated in all or two of the three L2/3 types in DR (**Figure 4F-H**). Conversely, most type-identity genes (87%; n=248/286) were vision-independent, consistently marking types A, B, and C in NR and DR (**Figure 4F,G**). Of 286 type-specific genes 38 (13%) were vision-dependent (**Figure 4F,I**). These *vision-dependent type-identity* genes fell into six groups based on their patterns of regulation (**Figure 4I**). Overall, type A had smaller fold changes in DR compared with types B and C in almost all genes. Unlike types B and C-specific genes, a group of type A genes (Group 2; n=9) were up-regulated in types B and C, but did not change in type A themselves. The changes in individual type-identity genes were consistent with the overall trend that type A was less affected by DR than types B and C.

Together, these results exemplify the plasticity of the transcriptional programs defining A, B and C cell types, resulting in a redistribution of cells in the 2D manifold associated with type identity due to visual deprivation (**Figure 4J**). The nature of this redistribution is consistent across biological replicates of snRNA-seq (**Figure S11**) and MERFISH (**Figure S12**). Finally, L2/3 glutamatergic neurons (also known as L2/3 intra-telencephalic neurons; L2/3 IT) are more sensitive to vision than deeper-layer intra-telencephalic (L5/6 IT) neurons, consistent with prior results (9) (**Figure S14**).

## Discussion

We previously showed that vision deprivation selectively impacts L2/3 glutamatergic neuronal types in V1 (9). Using unsupervised clustering of snRNA-seq data, we found that L2/3 neuronal clusters in DR mice “*poorly resembled the three types in NR animals, and the expression patterns of cell-type-specific marker genes were disrupted*”. We interpreted these results as “*a global disruption of gene expression patterns throughout L2/3*”, resulting in “*the loss of [L2/3] cell type identity in animals deprived of light*”. The nature of the disruption in gene expression was, however, not clear due to our inability to relate cell types between the two conditions.

Here, combining spatial transcriptomics with new computational analyses enabled us to relate L2/3 cell types between NR and DR and thereby clarify the vision-dependent gene expression changes. Visual deprivation impacts two orthogonal gene programs, one associated with experience-induced cell states and the other overlapping with programs of cell type identity. The deprivation-induced state changes dominate the transcriptomic variation in DR, masking cell type distinctions in the unsupervised clustering analysis used previously.

Using multi-tasking theory, we showed that the transcriptomic variation among L2/3 cells can be represented as a continuous manifold in 2D bounded by a triangle whose vertices represent archetypes A, B, and C. These archetypes represent the “basis sets” used to construct the profiles of sampled L2/3 neurons. While preserving the archetypes themselves, dark-rearing redistributes cells within the manifold, shifting cells away from archetypes B and C. Thus, L2/3 cell types are not lost in animals deprived of light, but their composition is altered, and the underlying gene expression programs are masked by orthogonal activity-induced changes. These changes are consistently reproduced in both MERFISH and snRNA-seq datasets.

A key tenet of multi-tasking theory is to link transcriptomic continua to the spatial zonation of functionally distinct cell types. Previous observations indicate that neurons within different L2/3 sublayers (A, B, C) differ in their functional properties and projections to higher visual areas (9, 11, 29). Our results support the idea that vision is required to maintain the proper distribution of cells in the L2/3 cell-type continuum and that vision preferentially impacts types B and C more than type A. The unequal dependence on vision could be related to the spatial zonation, as previous studies have shown that deep, but not more superficial, L2/3 cells receive LGN inputs (30). Consistent with this, our GO analyses suggest that types B and C but not type A genes are enriched in biological processes associated with sensory input (**Table S3**).

Previous studies have shown that neural activity influences the transcriptomes of the developing neurons (9, 11, 13, 31, 32), with L2/3 glutamatergic neurons being the cortical cell types that are most sensitive to sensory experience (13, 32), which is in agreement with functional studies (33, 34). Several studies have highlighted the regulation of IEGs under sensory deprivation, although the direction of regulation may depend upon the experimental protocol (9, 11, 32). Our data, based on both snRNA-seq and MERFISH, suggests dark-rearing itself leads to a massive up-regulation of IEGs and the effect is specific to visual areas. This area-specificity suggests the upregulation of IEGs is unlikely to be an experimental artifact. In addition, known IEGs only represent about 10% of genes altered by dark-rearing (**Figure 4F**).

How the interplay between intrinsic and vision-dependent molecular programs establish the transcriptomic continuum of L2/3 glutamatergic neurons, and how this sculpts circuitry and visual perception are exciting and challenging questions. Emerging techniques correlating transcriptomics, neural activity, and/or connectivity at the single neuron level provide new tools to explore these mechanisms (35–38). We anticipate that these studies, alongside modeling and computational analysis, will provide insights into the significance of the continuous nature of cell type variation for the proper function of V1. Multitasking theory (10, 39) suggests that transcriptomic continua aid the division of labor among cells in a tissue responsible for multiple functions. For continua bound by a polygon, cells close to each vertex (archetype) are specialized for a particular function, while cells in the middle are “generalists”, which can perform multiple functions at the cost of being sub-optimal for any single function. When gradients in gene expression space are also correlated with position, this can create spatial domains within a tissue, each suited for a unique complement of functions. Consistent with this idea, transcriptomic continua have recently been found to be a common trait in the mammalian brain, and they are often correlated with spatial and physiological continua (40–47). Indeed, the physiological and morphological features of L2/3 neurons in binocular V1 vary as a continuum along the pial-ventricular axis (48). Together, these findings suggest that vision controls the function of L2/3 glutamatergic neurons in V1, at least in part, by contributing to shaping the properties of this continuum. As L2/3 neuron types have selectively expanded in the primate cortex, continua of cell types may be particularly well suited to change by experience and may contribute to the enhanced cognitive capacities of primates.

## Supplementary Figures

**Figure S1.**
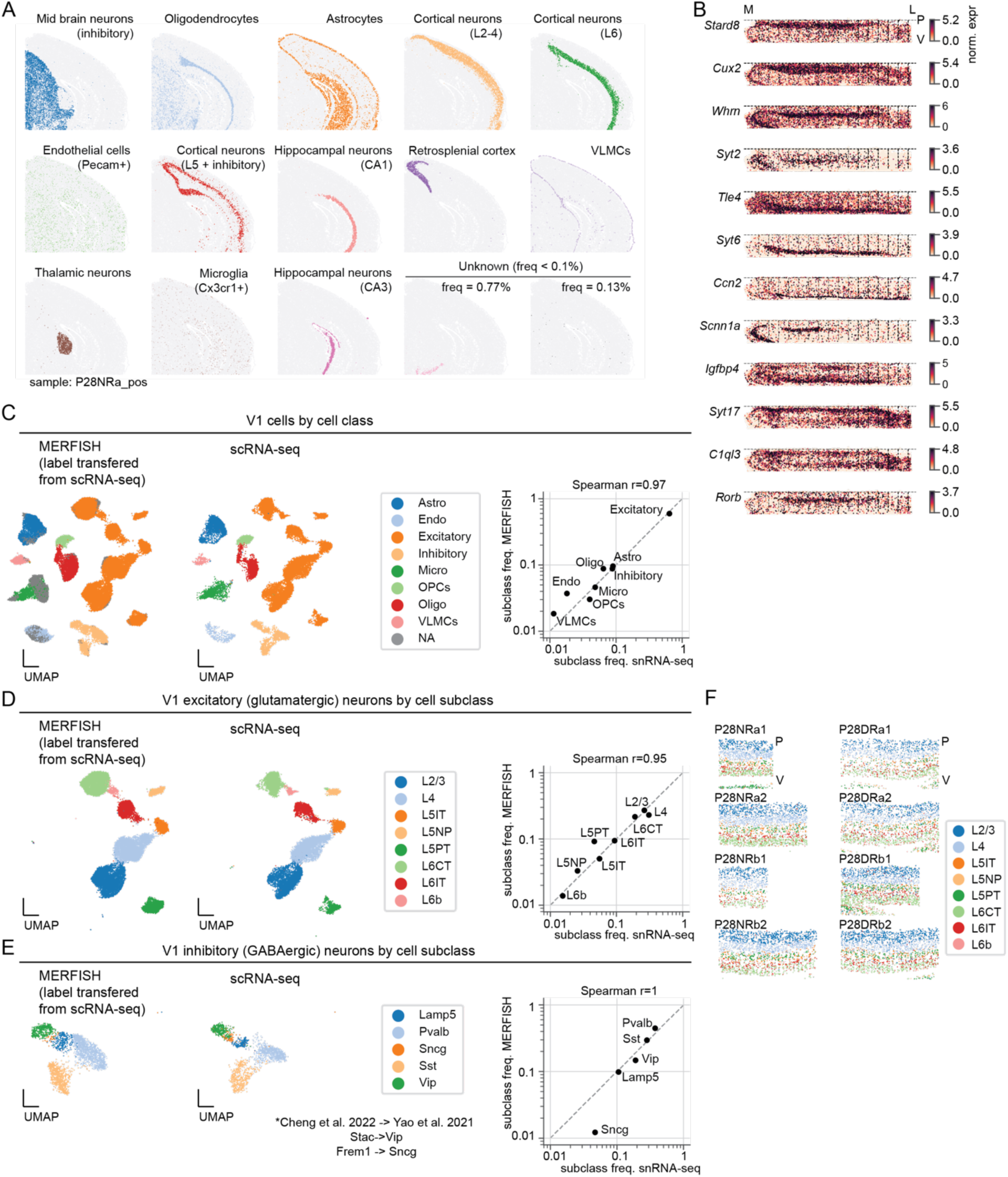
Related to Figure 1. MERFISH identifies major anatomical structures of the brain and cell subclasses in V1. (A) Spatial distributions of individual cell clusters identified based on gene expressions and spatial proximity (Related to Figure 1B). (B) *In situ* gene expression patterns across the straightened cortex. (C) V1 cell classes represented in UMAP obtained from integrating MERFISH (left panel; this study) and snRNA-seq (middle panel (9)) using Harmony (14). Right panel compares cell type proportions between snRNA-seq and MERFISH. 19.6% of MERFISH cells could not be classified (marked as NA and colored gray) due to their low quality and were filtered from downstream analyses. These cells are likely low-quality glia as they are co-clustered with non-neuronal cells. (D-E) Subclasses of V1 glutamatergic neurons (D) and GABAergic neurons (E) represented in UMAP embeddings obtained from integrating MERFISH (left panel; this study) and snRNA-seq (middle panel (9)) using Harmony (14). Right panels compare cell type proportions between snRNA-seq and MERFISH. (F) Spatial distributions of V1 glutamatergic subclasses.

**Figure S2.**
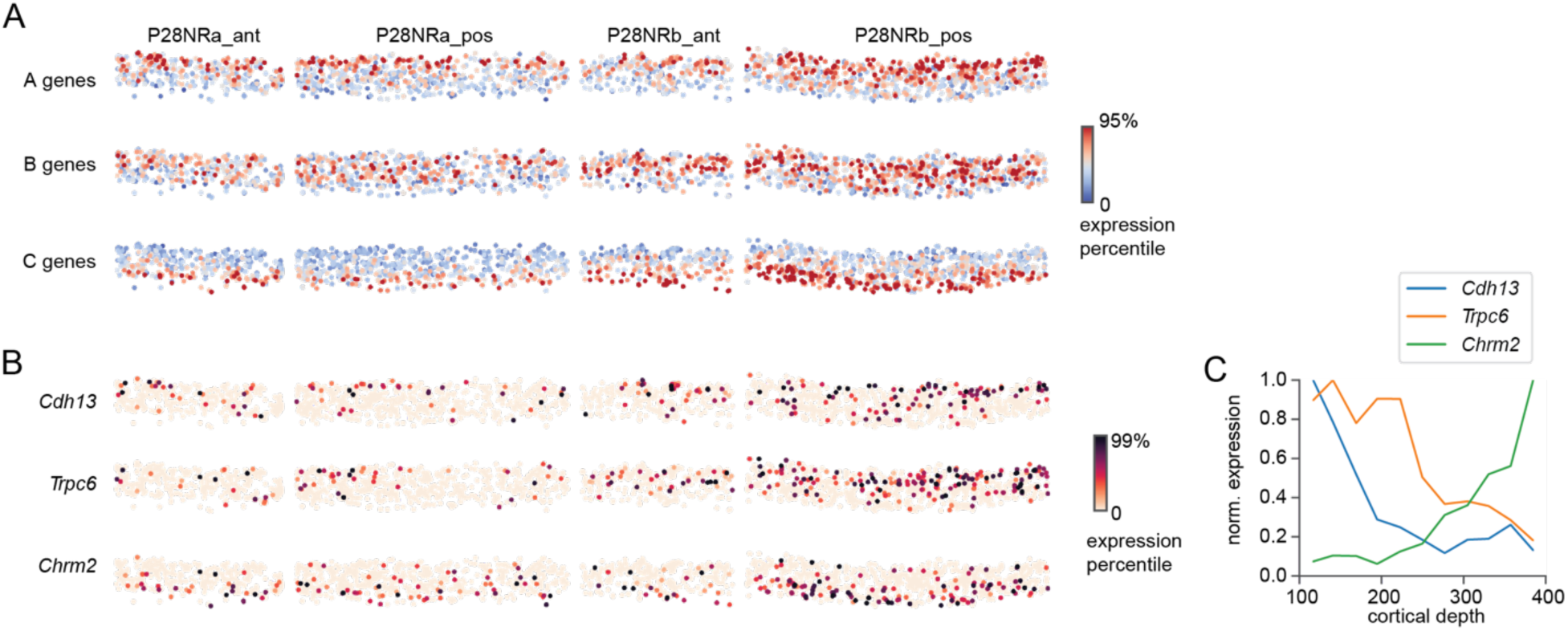
Related to Figure 1. *In situ* distributions of L2/3 type-identity genes. (A) *In situ* expression patterns of types A, B and C identity genes. (B) *In situ* expression patterns of *Cdh13*, *Trpc6* and *Chrm2*. *Cdh13* marks type A, *Trpc6* marks type B, and *Chrm2* marks type C. (C) Line plots showing the distribution of marker gene expressions along the cortical depths spanning L2/3.

**Figure S3.**
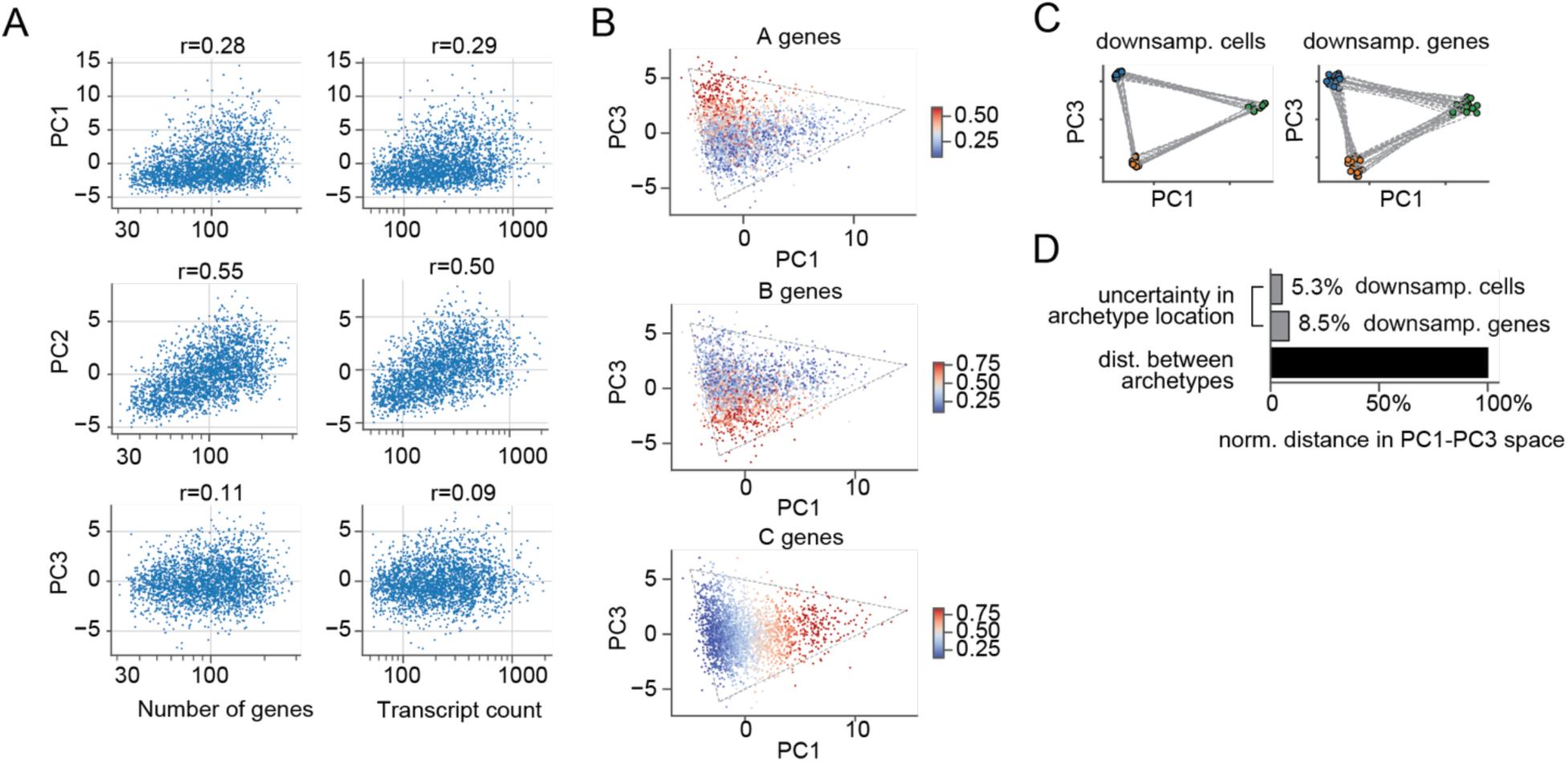
Related to Figure 2. Low dimensional representation of L2/3 cell types in MERFISH and inference of the bounding triangle from MERFISH data alone. (A) Scatter plots of PC1 (upper), PC2 (middle) and PC3 (lower) vs. the number of detected genes (left) and transcript count (right panel), respectively. PC1 is calculated using the 170 type-identity genes measured by MERFISH. PC2 correlates with these technical variables much more than PC1 and PC3. (B) V1 L2/3 cells embedded in PC1 and PC3. Cells are colored by the mean z-scored expression levels of type-A genes (upper panel), type B genes (middle panel) and type C genes (lower panel), respectively. (C-D) Stability of the bounding triangle. The bound triangle was inferred in several instances involving random down-sampling to 80% of cells (C, left panel) and 80% of genes (C, right panel). The stability of archetype locations are quantified by the % variation in archetype coordinates relative to the typical inter-archetype distances in these trials. Distances are measured by Euclidean distance in PC1-PC3 space. These inferences were done on MERFISH data, recapitulating results from snRNA-seq showed in Figure 2.

**Figure S4.**
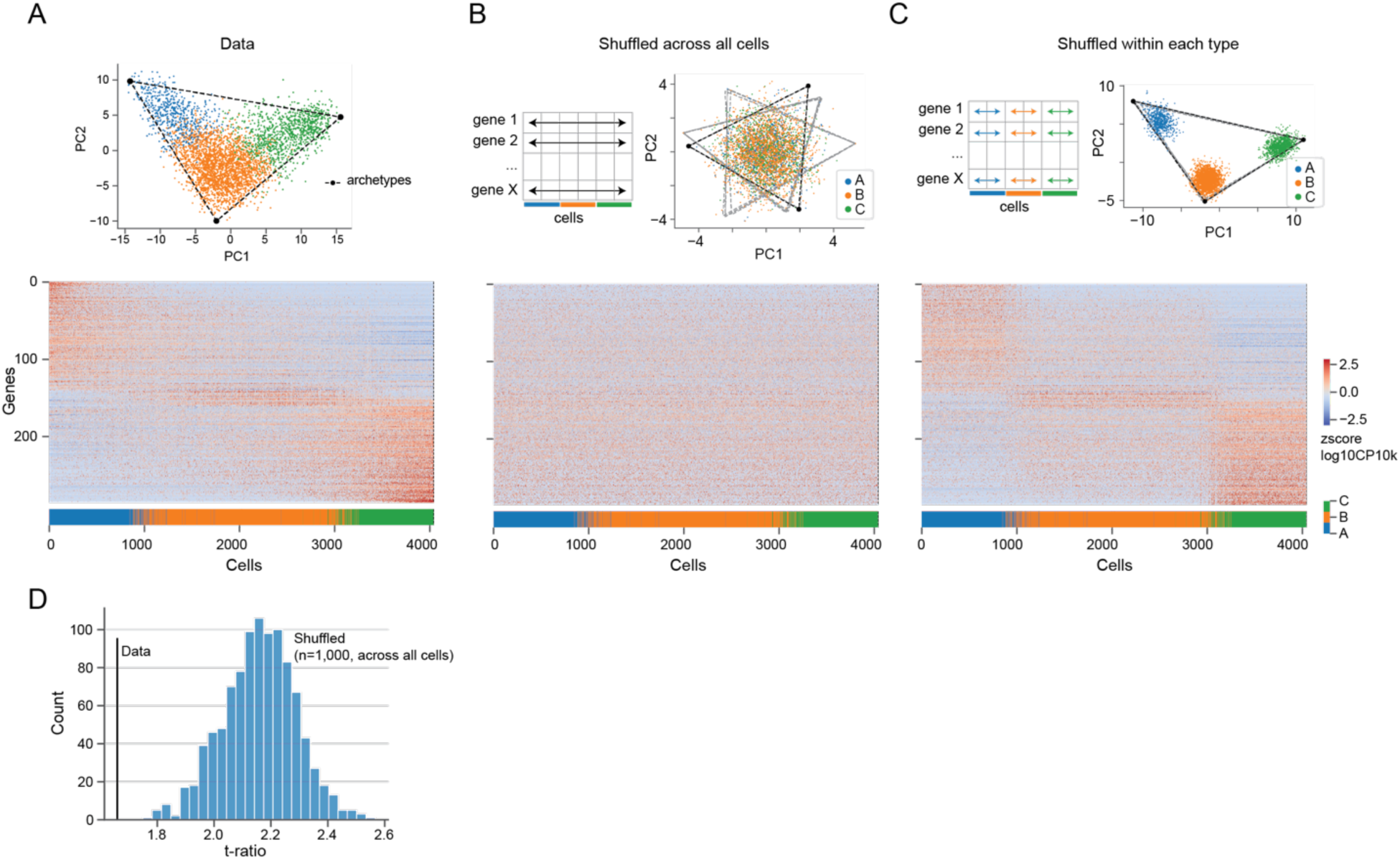
Related to Figure 2. Data shuffling disrupted the geometry of the L2/3 continuous transcriptomic manifold. (A) *Upper panel:* PCA embedding of L2/3 glutamatergic neurons in snRNA-seq using 286 type-identity genes. Cells were colored by type assignment based on the original clustering (9). The bounding triangle is inferred from archetypal analysis. *Lower panel:* expression profiles of L2/3 type-identity genes across all L2/3 glutamatergic neurons (n=4,044; P28NR). Expressions were quantified as z-scored, log- and size-normalized counts from scRNA-seq. Cells were ranked by diffusion pseudo-time (DPT; (19, 20)), and colored by type assignment. (B) Same as (A) but after shuffling each gene independently across all cells. (C) Same as (A) but after shuffling each gene independently across cells within each type. (D) Histogram of T-ratios for the data and shuffled data (n=1,000; shuffled across all cells). T-ratio is the ratio between the area of the convex hull and that of the principal convex hull (triangular fit).

**Figure S5.**
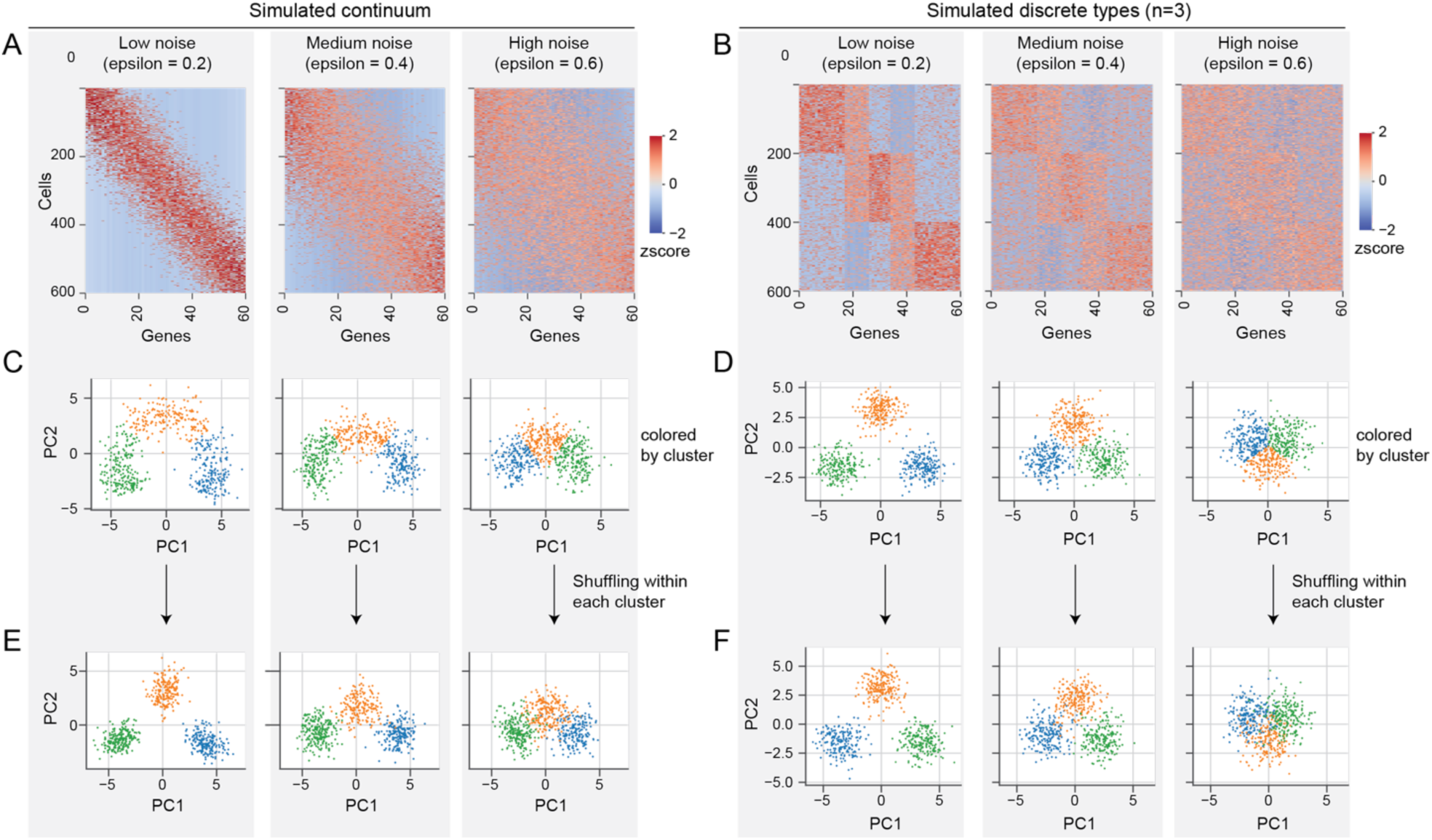
Related to Figure 2. A data shuffling procedure that split a continuum while preserving discrete types. (A-B) Expression profiles (gene by cell) for simulated continua (A) and simulated discrete types (B) each with varying degrees of noise level (parameterized by epsilon; see **Methods**). Expression was quantified as z-scores. (C-D) PCA embeddings (PC1 and PC2) of simulated continua (C) and simulated discrete types (D). Cells were colored by type. (E-F) Same as (C-D) but after shuffling each gene independently across all cells within each type. Shuffling genes within a cluster splits a continuum into separate clusters when noise is low (epsilon = 0.2 ∼ 0.4). The same procedure has no effect on discrete types whatsoever.

**Figure S6.**
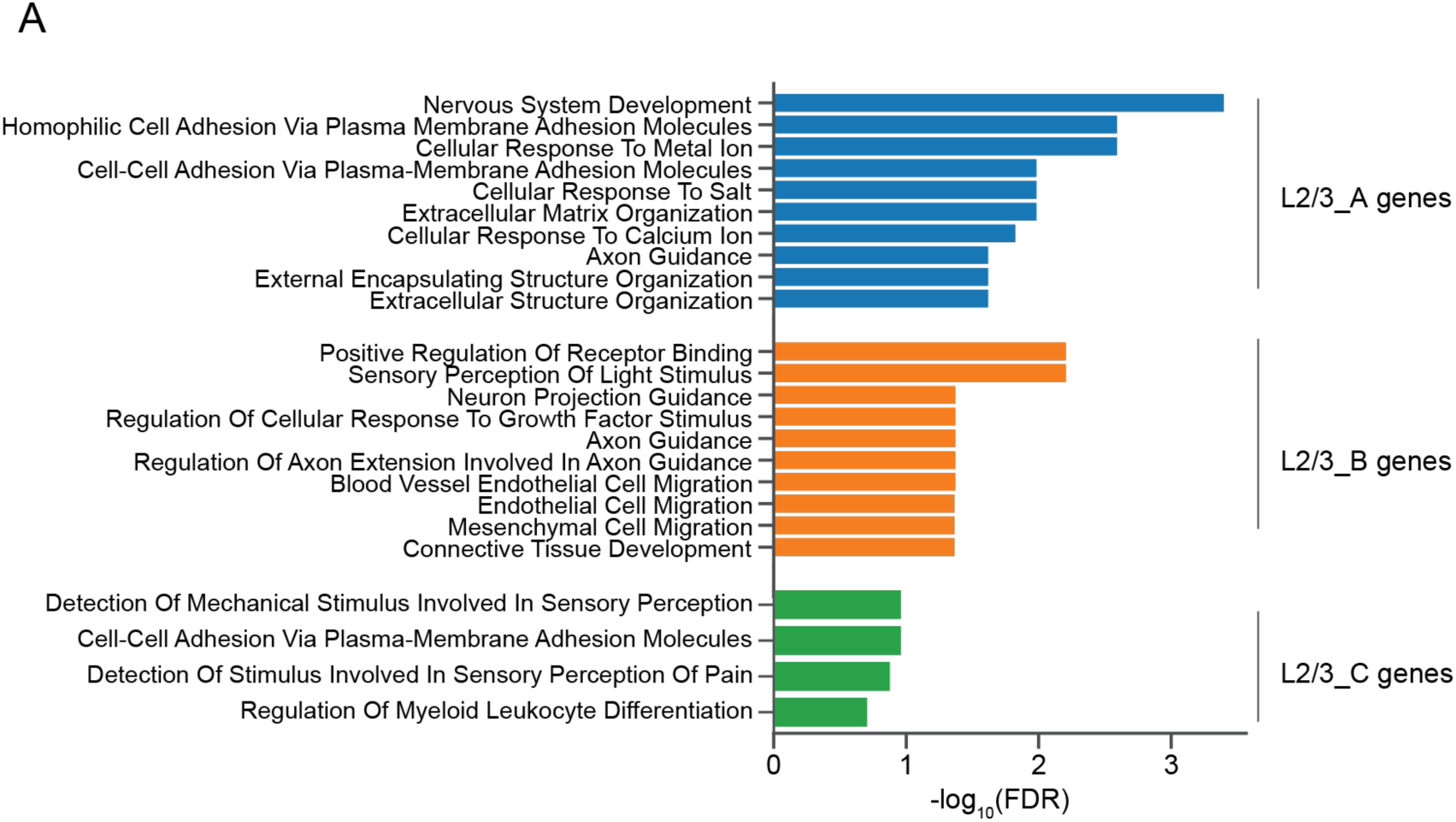
Related to Figure 2. Gene Ontology analysis identify enriched biological processes for type-identity genes. (A) Enriched biological processes for types A, B and C identity genes, respectively.

**Figure S7.**
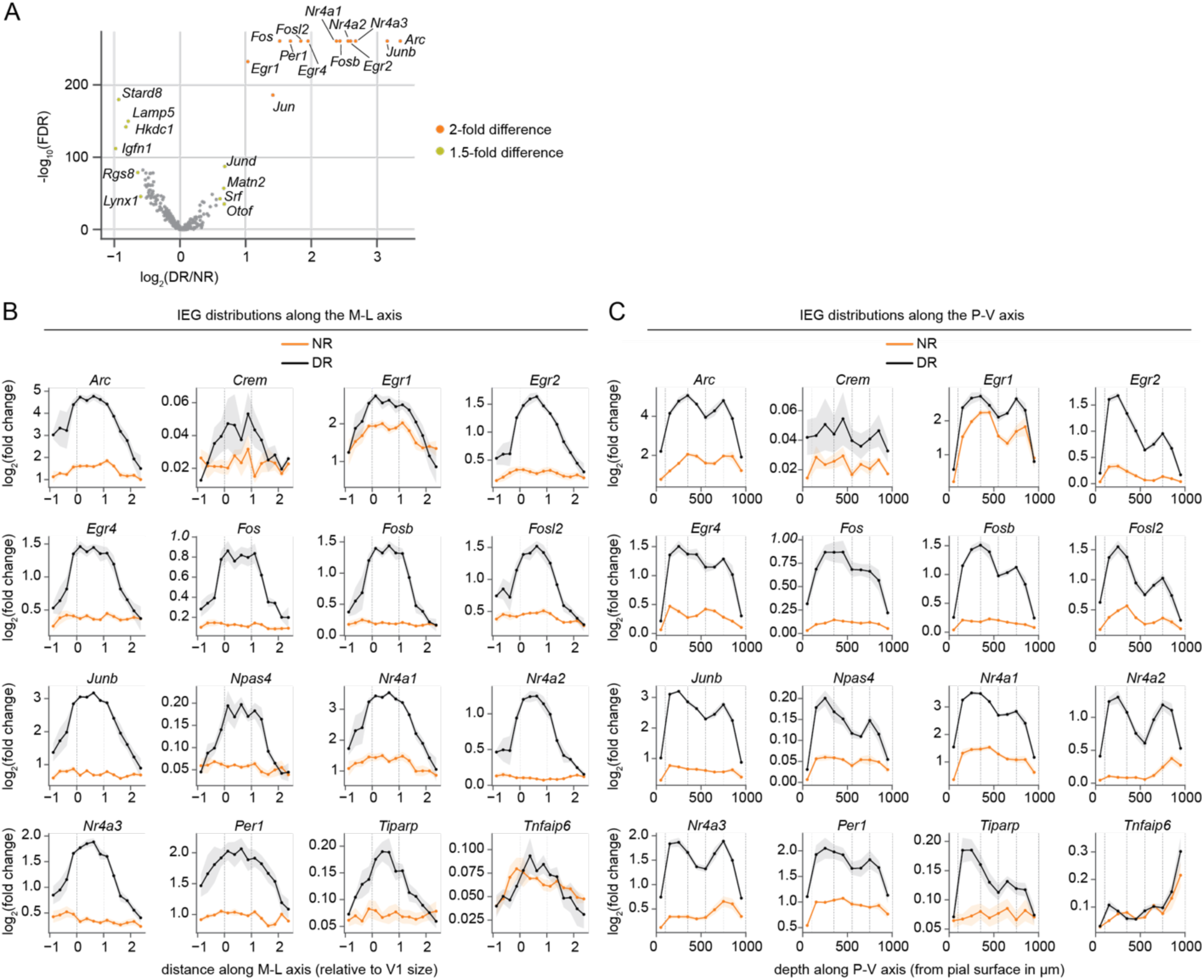
Related to Figure 3. Vision-dependent genes profiled by MERFISH. (A) Violin plot showing differentially expressed genes between NR and DR L2/3 glutamatergic neurons profiled by MERFISH. (B-C) Mean expression levels of individual IEGs along the M-L axis (B) with V1 in the middle and its flanking regions on the two sides, and along the P-V axis (C) across cortical layers. Lines demarcate V1 (B) and cortical layers (C).

**Figure S8.**
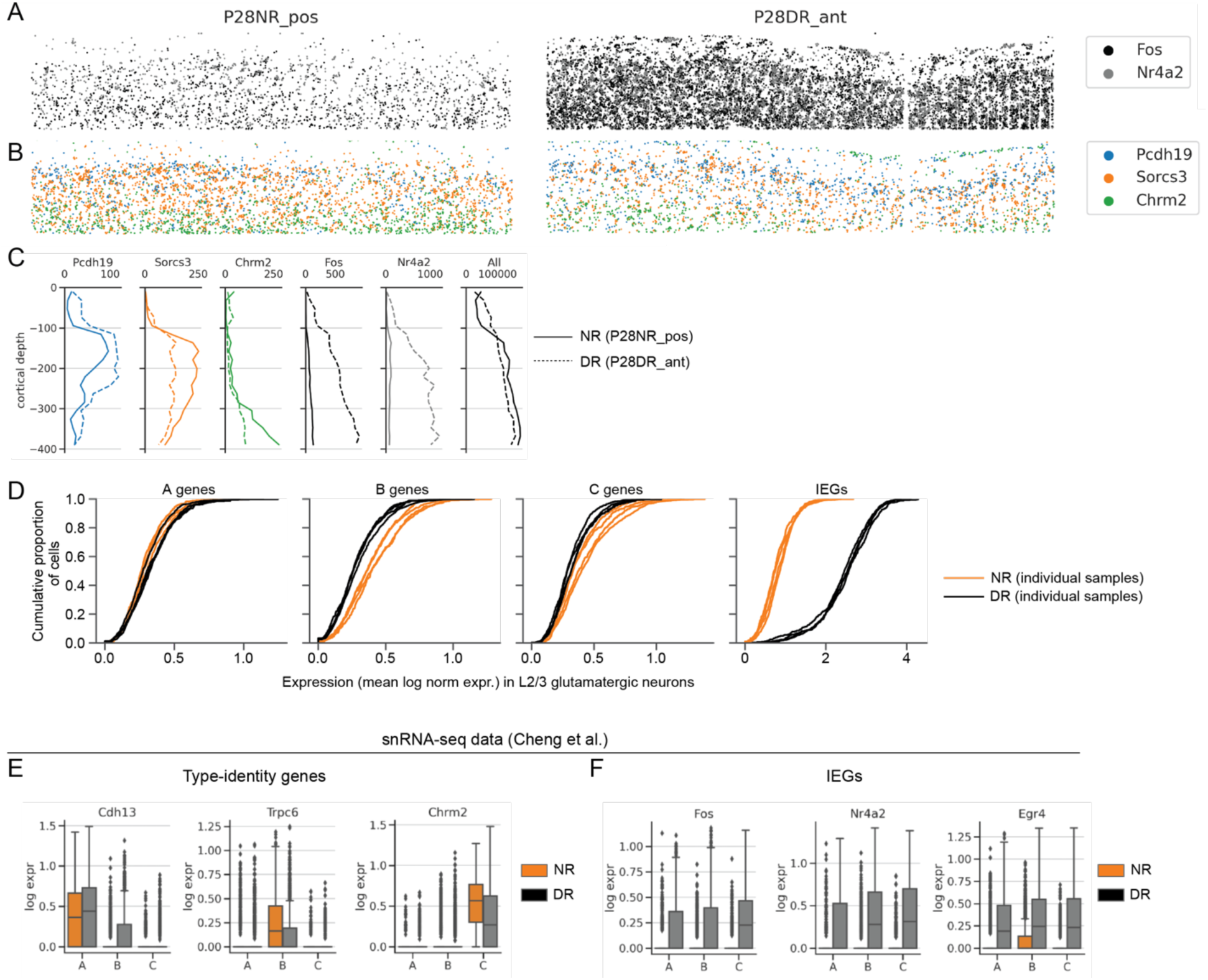
Related to Figure 3. MERFISH detected transcripts in V1 L2/3 for specific genes and gene groups. (A-B) Detected transcripts of Fos and Nr4a2 (A) and of *Pcdh19*, *Sorcs3*, and *Chrm2* (B) in V1 L2/3 for NR and DR samples. The entire V1 L1-3 area is shown. (C) Quantification of the number of detected transcripts at different cortical depths for genes shown in panels (A-B). (D) Cumulative distribution of gene expression across different gene groups and samples. Samples are colored by NR and DR. Group-level expression is defined as the mean expression level across genes within the group. (E-F) Gene expression distributions (in box plots) of selected type-identity genes (E) and IEGs (F) in L2/3 types from NR and DR mice.

**Figure S9.**
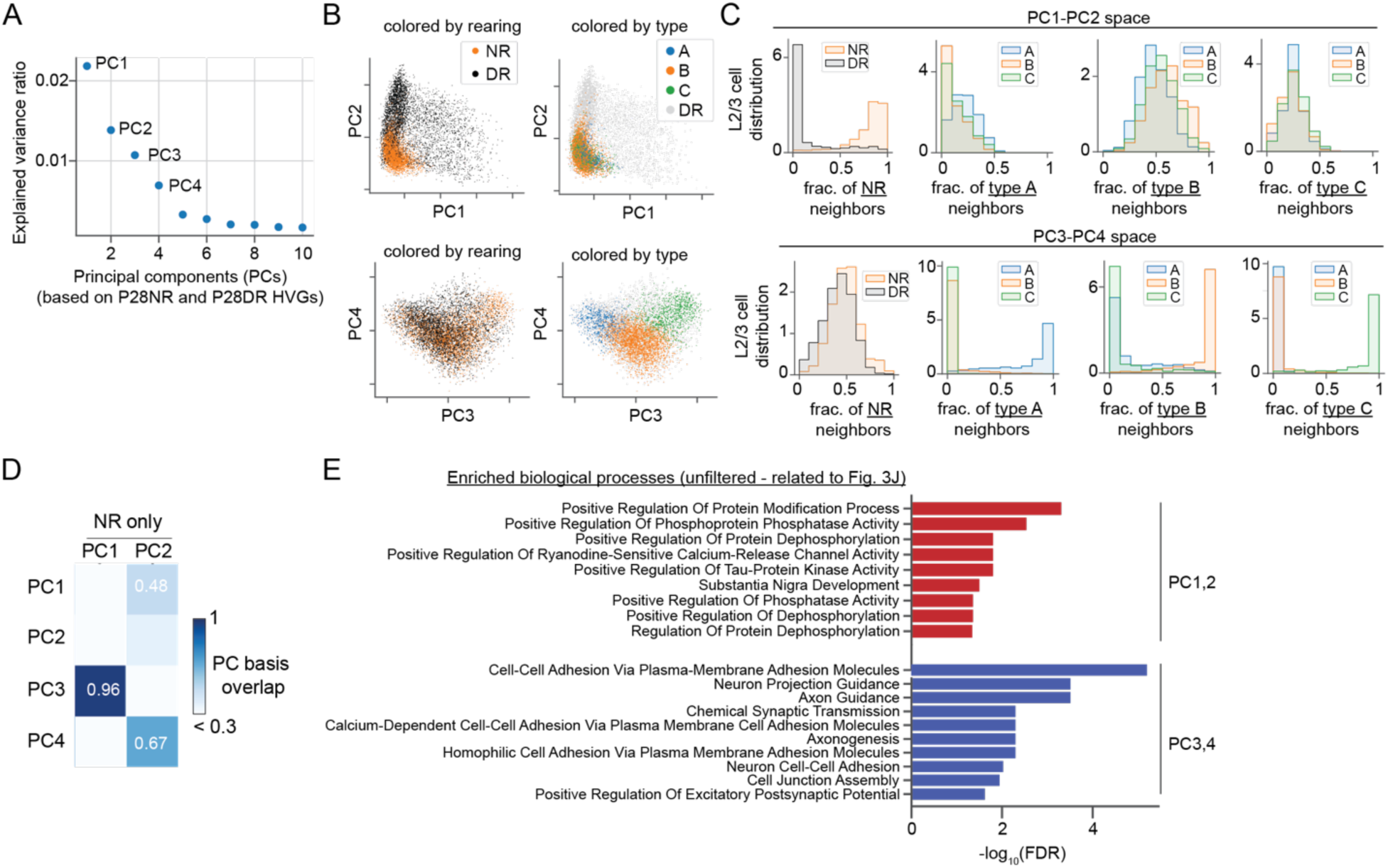
Related to Figure 3. PCA analysis revealed orthogonal components of transcriptomic variations in L2/3 transcriptomes from NR and DR mice. (A) Fraction of total variance captured within the top 10 principal components (PCs) in NR and DR (P28) L2/3 cells. (B) Distribution of NR and DR cells in PC1-PC2 space (upper) and PC3-PC4 space (lower). PCs are computed from n=6,360 HVGs. Cells are colored by rearing condition (NR vs DR) (left) and by cell type (A, B and C) for NR cells (right). (C) Distribution of neighbor identities. Column 1 shows the fraction of NR neighbors for NR cells (in orange) and DR cells (in black) in PC1-PC2 space (upper panel) and PC3-PC4 space (lower panel). Columns 2-4 show the fraction of types A, B and C neighbors respectively. (D) Pairwise overlap between PCs derived from using both NR and DR cells versus using NR cells only. (E) Enriched biological processes for PC1-PC2 and PC3-PC4 driving genes. This is related to main Figure 3J.

**Figure S10.**
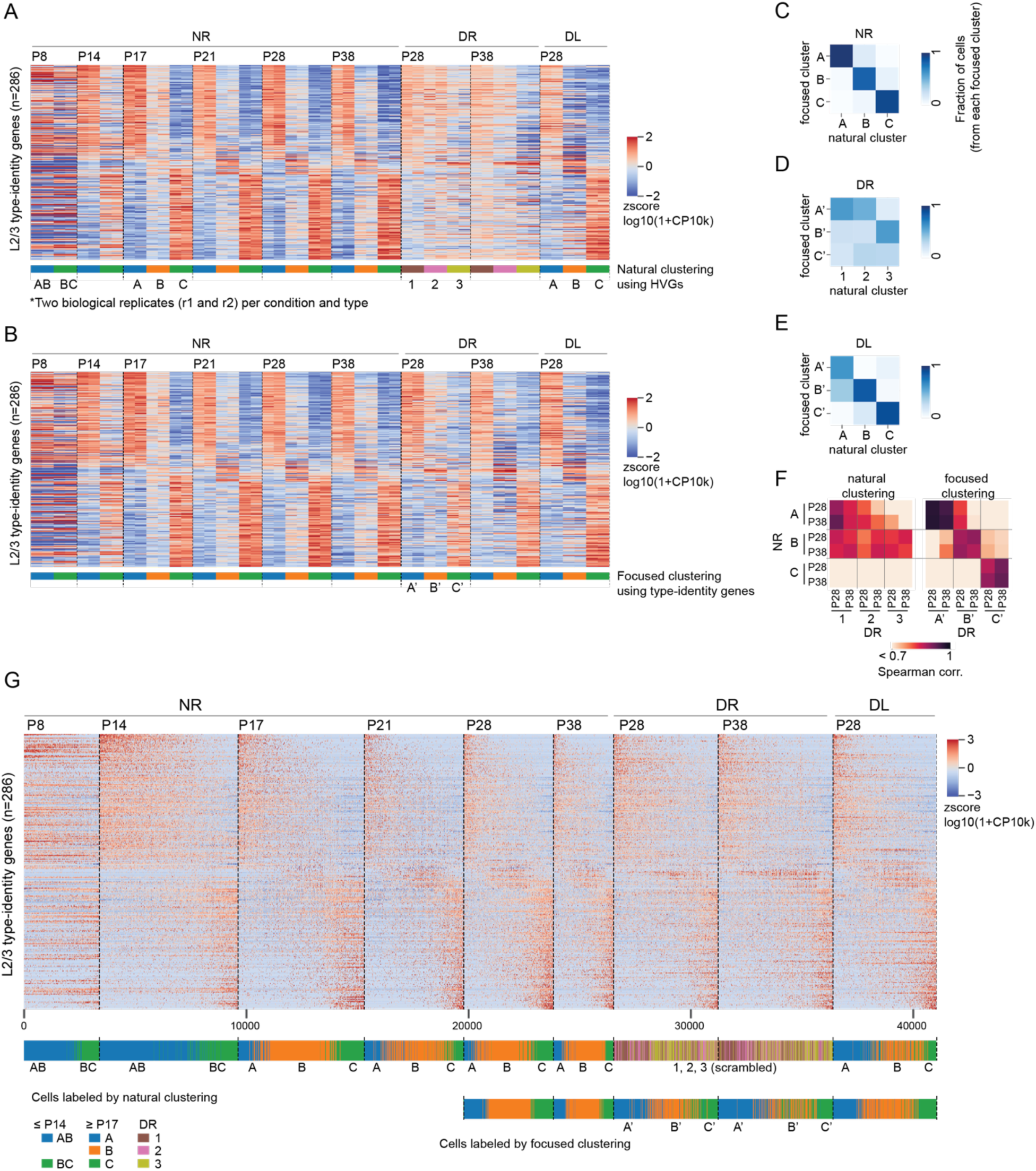
Related to Figure 3. The L2/3 transcriptomic continuum across time and conditions. (A-B) Expression profiles of L2/3 type-identity genes for clusters based on natural clustering using HVGs (A), and based on focused clustering using type-identity genes (B). Natural clustering using HVGs mask the cell-type signatures in DR, while focused clustering using type-identity genes recover the cell-type signatures. (C-E) Confusion matrix of natural clustering vs focused clustering for NR (C), DR (D) and DL (E), respectively. (F) Pairwise Spearman’s correlation coefficients between NR and DR cell clusters based on L2/3 identity genes. Focused clustering has 1:1 correspondence between NR and DR clusters, while natural clustering does not. (G) Expression profiles of L2/3 type-specific genes for individual cells. Top: Gene expression profiles. Bottom: cells colored by HVG-based and focused clustering respectively. Cells were ordered by diffusion pseudo-time for each condition.

**Figure S11.**
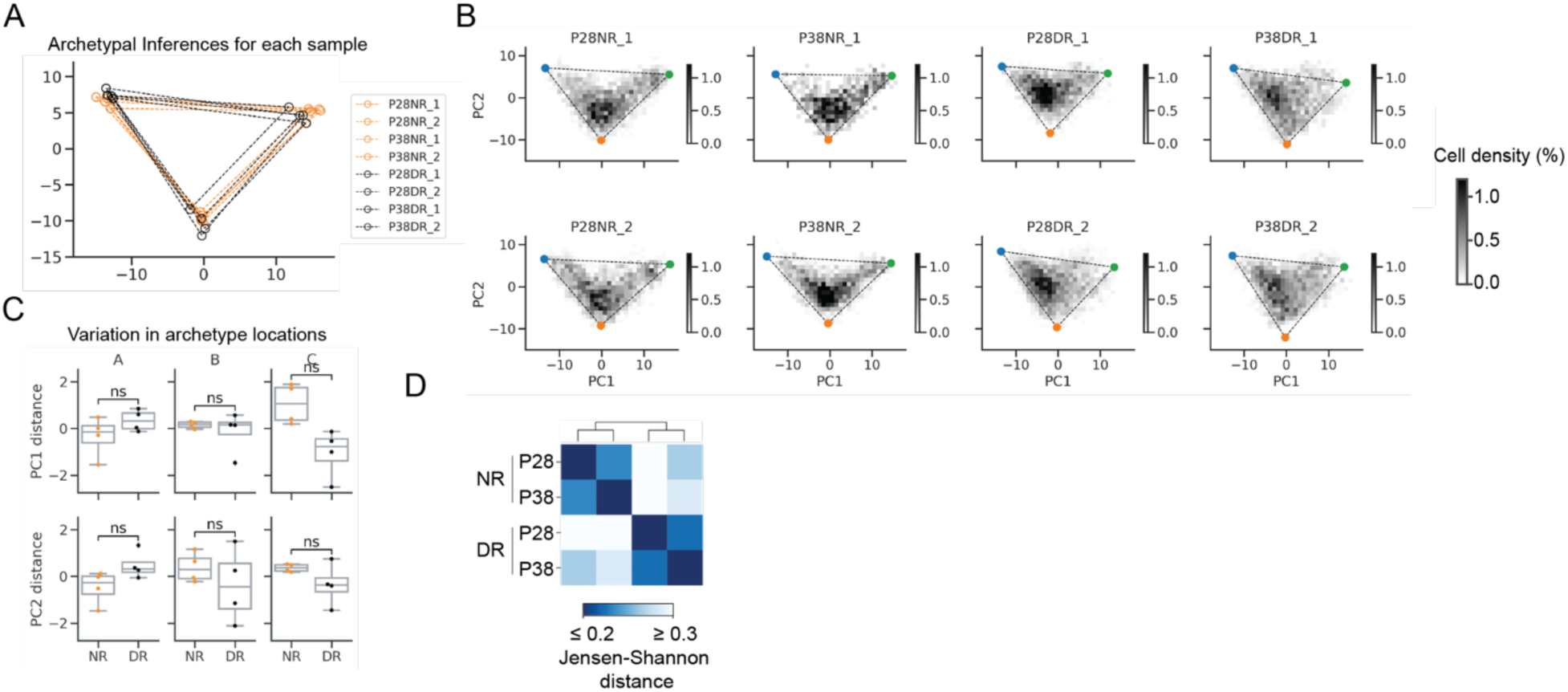
Related to Figure 4. Dark-rearing shifted cells along the continuous transcriptomic manifold of L2/3 cell types (snRNA-seq data). (A) Inferred L2/3 transcriptomic triangle for individual biological samples. (B) L2/3 cell density plot in PCA embeddings (PC1 and PC2 based on type-identity genes). L2/3 cells from all samples are embedded into the same PC1-PC2 space using type-identity genes. Cell density and the archetypal inference are done separately for each sample. (C) Boxplots showing the variations of archetype locations in PC1-PC2 space. (D) Pairwise Jensen-Shannon distances between samples of different rearing conditions.

**Figure S12.**
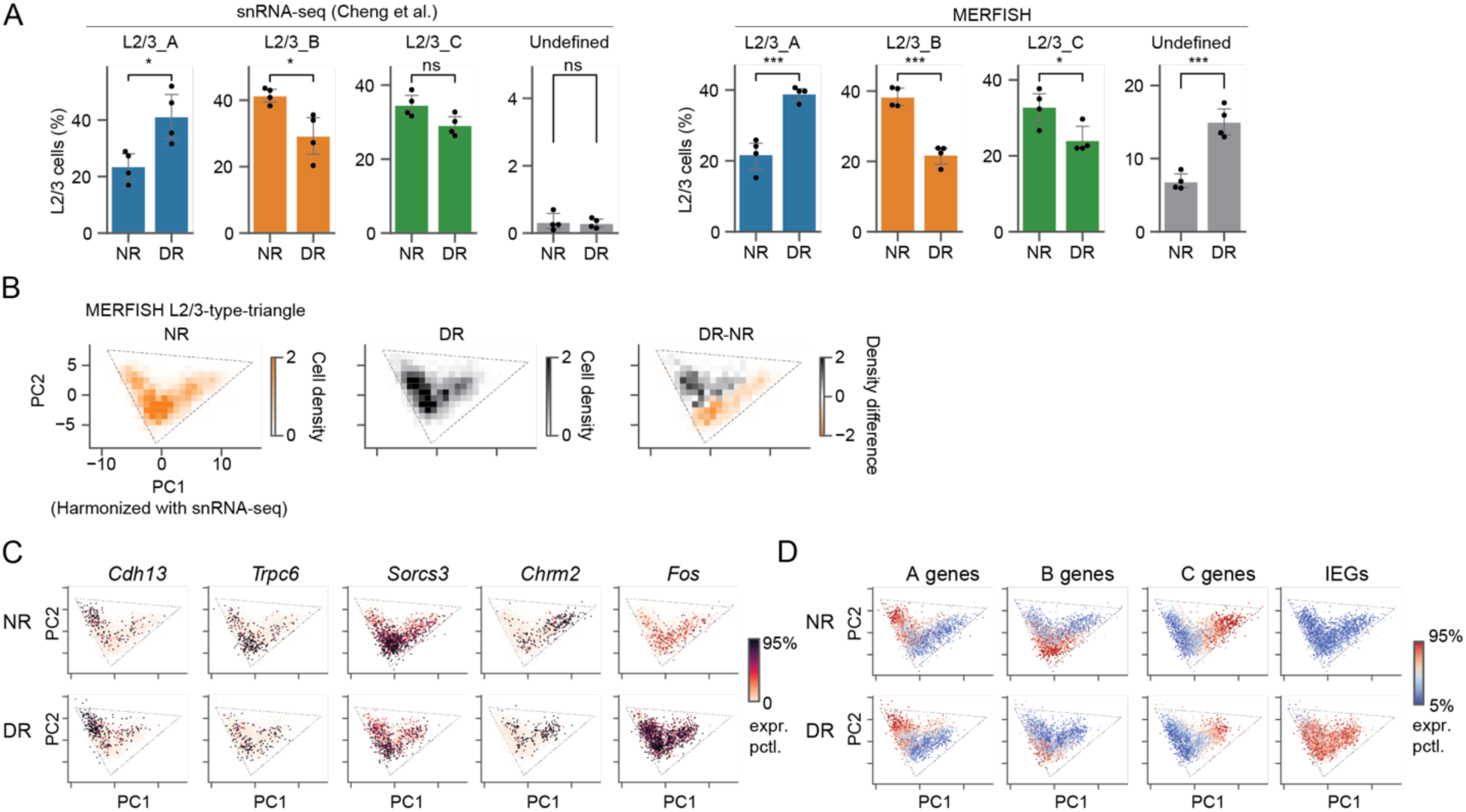
Related to Figure 4. Dark-rearing shifted cells along the transcriptomic manifold of L2/3 cell types (MERFISH data). (A) Proportion of L2/3 cell types from NR and DR samples profiled by snRNA-seq (left panel) and MERFISH (right panel). Each dot represents a biological sample. Cell type labels are based on the expression levels of type-identity genes (see Methods). (B) Distribution of L2/3 cells within its bounding triangle for NR and DR mice profiled by MERFISH. (C-D) Gene expression patterns for individual genes (C) and groups of genes (D) on the L2/3 type-triangle.

**Figure S13.**
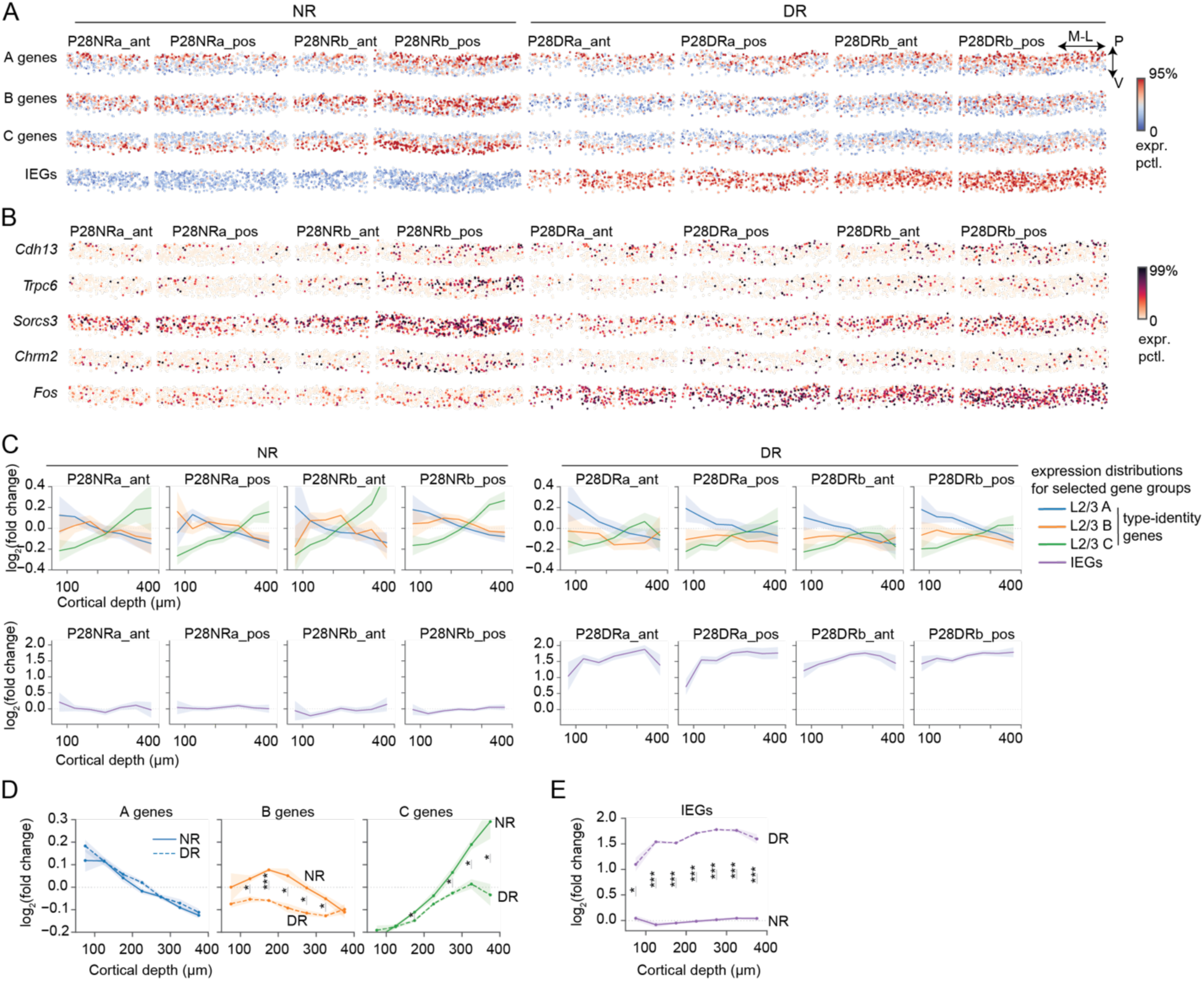
Related to Figure 4. Characterization of *in situ* gene expression in V1 L2/3. (A-B) in situ expression patterns of different gene groups (A) and individual genes (B) in V1 L2/3. Each row shows one gene (or gene group) and each column one sample. (C) Line plots showing the mean expression levels of type-identity genes (upper panel) and IEGs (lower panel) along the cortical depth for individual samples. (D-E) Line plots comparing the expression levels of type-identity genes (D) and IEGs (E) in NR and DR across cortical depths spanning L2/3. FDR < 0.05 (*) and FDR < 0.001 (***).

**Figure S14.**
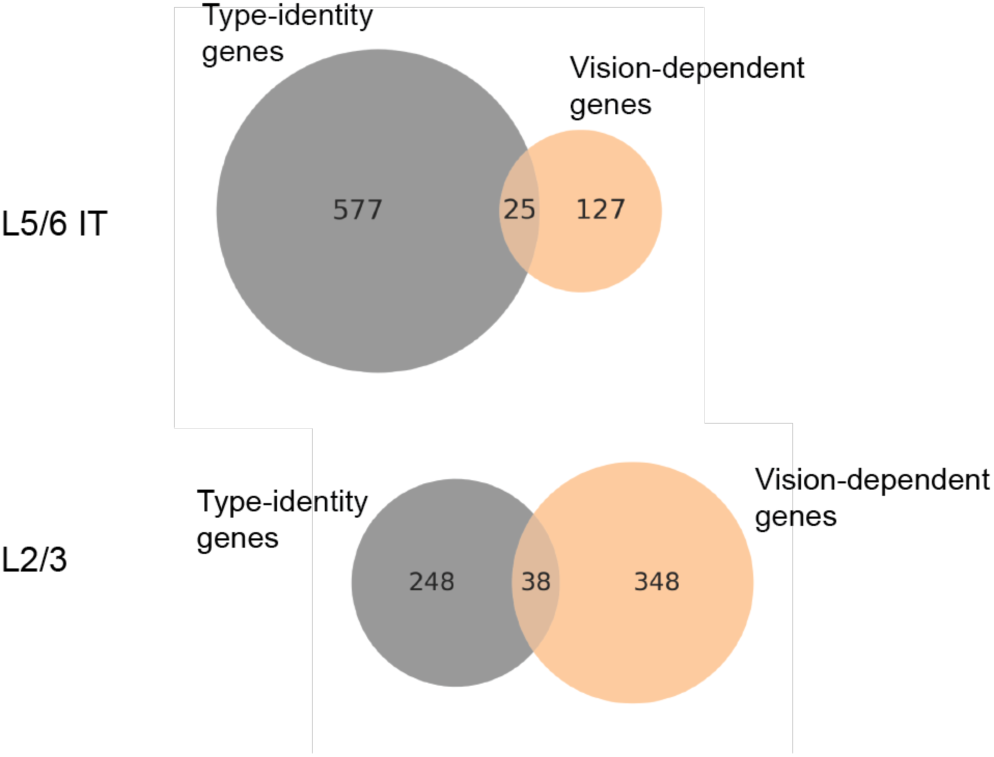
Related to Figure 4. Comparing type-identity gene programs and vision-dependent gene programs in superficial (L2/3) and deep (L5/6) layer intra-telencephalic (IT) neurons. Venn diagrams showing the numbers and the degree of overlaps between type-identity genes and vision-dependent genes identified in L5/6 IT (upper panel) and L2/3 IT (lower panel) neurons.

## Supplemental Tables

**Table S1.** L2/3 type identity genes (n=286), in which 38 were up- or down-regulated in dark reared mice.

**Table S2.** MERFISH gene panels (n=500).

**Table S3.** Enriched GO biological processes in L2/3 type-identity genes.

**Table S4.** Enriched GO biological processes in PC1-PC2 and PC3-PC4 top loading genes.

**Table S5.** Previously identified L2/3 stimulus-response genes (from ref (11)).

## Methods

### Mice for MERFISH

Mouse breeding and husbandry procedures were carried out in accordance with the animal care and use committee protocol number 2009-031-31A at the University of California, Los Angeles. Mice were given food and water *ad libitum* and lived in a 12-hr day/night cycle with up to four adult animals per cage. Only virgin male C57BL/6J wild-type mice were used for MERFISH experiments to match with Ref (9).

### Visual deprivation paradigm

Mice were dark-reared starting postnatal day 21 (P21), which marks the beginning of the ocular dominance critical period. Prior to P21, mice were reared in cages in a room with a 12-hour light/dark cycle. At P21, mouse cages were moved to a separate room and placed inside a dark box lined on the inside and the outside with black rubberized fabric (ThorLabs Cat# BK5) with edges sealed by tape and fabric to avoid any light entry. Mice were dark-reared in this setup for 7 days (P21-28DR group). During this period, any handling of the cages was performed in the dark with room lights off, door crevices sealed from exterior light, and in dim red light, which is invisible to the mice. A light meter was used to measure the amount of light inside the dark-rearing box before visual deprivation paradigm. All mice were dark-reared in 0-0.01 lux conditions. At the end of dark-rearing, mice were euthanized and harvested for MERFISH experiments under red light.

Normally reared mice were housed in a 12hr light-ON, 12hr light-OFF cycle and were harvested during a range of 6-8 hours into the light-OFF phase. The cages of normally reared mice were wrapped with the same black rubberized fabric and carried to dark room with only red light on to minimize light exposure before euthanasia.

### Tissue preparation and MERFISH experiments

All mice were anesthetized with isoflurane and then perfused transcardially with 10mL heparinized PBS. Following perfusion, the brains were harvested, imbedded in pre-chilled Tissue-Tek^®^ OCT mounting medium, and flash-frozen in dry ice-prechilled methylbutane. The frozen blocks were then kept on dry ice and stored at -80℃ until time of sectioning. To prepare cryosections for MERFISH, two entire OCT blocks (two brains) or three bisected OCT blocks (three hemibrains) were combined and sectioned at -20 °C in a Leica CM1850 cryostat. Two 12-mm-thick coronal slices (one anterior and one posterior containing binocular V1 with ∼550um apart in the anterior-posterior axis) were directly collected onto a specially coated 4cm-diameter MERFISH glass slides (merslides, Vizgen Item# 10500001). For locating the visual cortex, atlas coordinates from (Franklin and Paxinos, 2012) and Allen Brain Atlas were used. Collected cryosections on the MERFISH slides (merslides) were fixed in 4% paraformaldehyde in PBS (15min at RT) in a 6cm petri dish, rinsed with cold RNase-free PBS, and stored in 70% ethanol at 4 °C until the step of MERFISH probe hybridization.

MERFISH was performed according to Vizgen’s instructions. Briefly, merslides with brain sections stored in 70% ethanol were rinsed with Vizgen Sample Preparation Buffer (SPB) after aspiration of 70% ethanol, incubated with Vizgen’s Formamide Wash Buffer (FWB, 30 min at 37 °C) followed by MERFISH probe labeling with a customized mouse gene panel containing 500 mouse genes (VZGCP0991, **Table S2**, ∼40h at 37 °C) in a moist chamber, and washed with FWB twice (30 min at 47 °C). The brain sections were be embedded with Vizgen gel mix after removal of FWB, cleared in 5mL clearing mix solution with 50 mL protease K (overnight at 37 °C), stained with Vizgen DAPI/poly(T) reagent included in the Vizgen 500-gene imaging kit (10 min at RT) after rinse with SPB and FWB (10 min at RT). Then the merslide was thoroughly rinsed with SPB, carefully assembled into an imaging gasket chamber, and uploaded into the MERSCOPE for imaging. The MERFISH imaging was done on the MERSCOPE with an activated Vizgen 500-gene imaging kit after adding the imaging buffer activator and RNase inhibitor (100 mL) as instructed. The imaging process was conducted under the direction of Vizgen’s MERSCOPE program (Software version 233.230615.567) with the default settings (both polyT and DAPI channels “on”, scan thickness: 10mm). Once the MERFISH imaging process was completed, the output files were transferred for in-depth analysis with MERFISH Visualizer and in-house custom bioinformatic pipeline.

### snRNA-seq data processing and normalization

Cell-by-gene count matrices from the previous study were downloaded from Gene Expression Omnibus (GEO) repository <u>GSE190940</u> (9). Cell type labels were downloaded from the associated Github repository https://github.com/shekharlab/mouseVC. Raw count matrices were normalized as described before. Transcript counts within each cell were rescaled to sum up to 10,000. A pseudo-count of 1 was added to the normalized transcript counts for each gene within each cell, followed by log10-transformation. For PCA and clustering, log10-transformed counts were z-scored across cells for each gene.

We reproduced a list of L2/3 type-identity genes (n=286, **Table S1**) by following the differential expression analysis used in the previous study. Briefly, the expression of each gene was compared in one type versus others (L2/3 A vs B-C; L2/3 B vs A-C; and L2/3 C vs A-B). We used *scanpy.tl.rank_genes_groups* to identify differentially expressed genes if a gene meets the following criteria: a) false discovery rate (FDR) < 0.05 based on Wilcoxon rank-sum test; b) fold change > 2, and c) the gene was expressed in > 30% cells in the up-regulated type.

### MERFISH data processing and normalization

Starting from a cell-by-gene matrix, in which each entry is an integer representing the number of transcripts detected in each cell and for a specific gene by MERFISH, we first removed low quality cells. We kept cells with a volume between 50∼2000, total number of detected transcripts > 10, and false positive rate of transcript detection <5% for the initial analysis. (Later to identify L2/3 cell types A, B and C, we further selected high-quality cells with total number of detected transcripts > 50). Notably, the false positive rate is a unique feature of MERFISH. In MERFISH experiments, transcripts were identified by binary barcodes extracted from repeated rounds of hybridization, imaging and puncta detection. A few barcodes were designed to be blank barcodes unmapped to any genes, which allowed us to estimate false positive transcript detection rate. Out of the eight MERFISH sections, four were done on the full coronal sections while the rest were done on hemibrain coronal sections only. For consistency, we computationally cut out a hemi-brain section for every sample for downstream analysis.

Following the same procedure employed in a previous MERFISH study (18), we normalized the raw count matrix first by cell volume and then by mean transcript count across samples such that each section had the same mean number of transcripts (n=250) detected per cell. This procedure effectively ameliorated cell volume-associated variation and batch effects across samples. After the above cell-to-cell and sample-to-sample size normalization, we normalized the data by log (+1) transformation and z-scored every gene.

Notably, we found a massive up-regulation of a panel of IEGs in sections from DR mice compared to NR (see main text). To remove this confounding factor, sample-to-sample normalization was done using 484 out of the 500 genes profiled by MERFISH (excluding the 16 IEGs) to compute the total transcript counts. We found that the identification of major cell subclasses was unaffected by the inclusion/exclusion of IEGs. However, IEG removal helped quantify gene expression changes in NR vs DR within L2/3 types more accurately.

### Spatial domain analysis

To identify the major anatomical areas and molecular organization within our MERFISH sections, we performed a clustering analysis on each section based on cell-to-cell gene expression similarity and spatial proximity. Using the normalized data as features, we performed a principal component analysis (PCA) and generated a gene-expression-similarity graph based on coordinates in the space of the top 50 principal components (PCs). We also generated a spatial-proximity graph based on spatial locations of the cells. Both graphs were generated using *scanpy.pp.neighbors* following default parameters. We generated a spatial-and-gene hybrid graph by blending the spatial and the gene graphs with equal weighting. We applied Leiden clustering on the hybrid graph using *scanpy.tl.leiden* following default parameters. The resulting spatial domains recover major anatomical structures in the mouse brain, which were visualized in **Figures 1** and **S1**. Clusters were annotated based on anatomical and molecular features.

### Establishing curved cortical coordinates and identifying V1

We developed an automated iterative procedure to fit a smooth curve along the pial surface, which served as a reference line to establish curved coordinate system along the vertical (pia-ventricular; P-V) and the tangential (medial-lateral; M-L) axis of the cortex.

We took advantage of meningeal cells (VLMCs), which specifically express *Slc6a13* and are located along the pial surface. Using these *Slc6a13+* cells as anchors, we first fit a fourth-order polynomial using *numpy.polyfit*. Subsequently, we used the fitted line as a reference to calculate the depth for all the anchor cells. Anchor cells whose depth that are >1/2 of the maximum depth (robust maximum taken as the 95% max depth) among all anchors were removed from the set of anchor cells for another round of more refined fitting. This procedure was repeated for 5∼10 times until the inferred reference line was stable and visually matched the cortical surface.

Using the reference line, all cortical cells were assigned a depth along the P-V axis and a width along the M-L axis.

Using the curved coordinates, cortical cells, defined as having cortical depth less than 1100 microns from the pial surface, were extracted from the full coronal sections for downstream analysis. V1 cells were identified based on tangential locations based on areal-specific markers mentioned in the main text.

### Identification of subclasses of V1 cells and integration with published snRNA-seq data

To systematically identify major cell populations, V1 cells were extracted and merged from all MERFISH sections (8 in total for this study; NR and DR) as mentioned above and were integrated with our previously published snRNA-seq data (9) using Harmony (14). We did this at two levels – the class level and the subclass level within the excitatory and inhibitory classes (**Figure S1**).

To integrate the MERFISH and snRNA-seq data (P28NR), we intersected MERFISH genes (n=484, excluding IEGs) and highly variable genes (HVGs; n=7,340) identified by snRNA-seq, which resulted in 361 overlapping genes between the two datasets. For this analysis, we only used high-quality MERFISH cells, defined as cells with at least 50 transcripts. We computed PCA (n=20 components) using cells from both datasets based on the overlapping genes (z-scored separately for each dataset). We then applied Harmony using *scanpy.external.pp.harmony_integrate* to generate harmonized PCs.

These harmonized PCs were used for downstream visualization and cell-type assignment. UMAP were generated using *scanpy.pl.umap* with the default parameters. Cell type labels were transferred from the published snRNA-seq data to MERFISH based on k-nearest-neighbor-based assignment using *sklearn.neighbors.NearstNeighbors*. A MERFISH cell was assigned to the most frequent cell class out of its k=100 snRNA-seq neighbors. To assign cell subclasses with a cell class, k=30 neighbors were used.

A small fraction of MEFISH cells could not be confidently assigned a cell class (**Figure S1C**). These cells were defined as those whose k=100 neighbors in the harmonized PCs contains ≤ 2 cells from snRNA-seq, indicating an unsuccessful data integration for these cells. These cells were assigned label “NA” (**Figure S1**), most likely being low-quality non-neuronal cells with lower number of transcripts, and unspecific spatial signatures. They were mostly grouped with microglia and endothelial cells in the UMAP embedding (**Figure S1C**).

### Identification of L2/3 glutamatergic cell types

The procedures above identified all cell subclasses, among which we focused on one subclass, the L2/3 glutamatergic neurons, for further analysis. Two strategies were used to further identify the three types of L2/3 neurons (types A, B and C). The first was to repeat the analysis above at the cell type level, which assigned each MERFISH cell an identity based on its k-nearest neighboring snRNA-seq cells in the harmonized PC space.

We also used a second strategy, which offers more biological interpretability. Given the continuous nature of L2/3 types bounded by the three archetypes A, B and C, we developed continuous A-B-C scores based on the expression levels of type-A, B and C identity genes (n=170 profiled by MERFISH). Each cell was assigned three scores (*p*_*a*_, *p*_*b*_, *p*_*c*_), with *p*_*a*_ + *p*_*b*_ + *p*_*c*_ = 1, representing the probabilities that this cell belongs to type A, B and C respectively. For a cell *i*, we first characterized its overall expression levels of type-A specific genes by the mean z-scored type-A-identity gene expression,

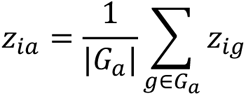

where *G*_*a*_ represents the set of type-A-identity genes and |*G*_*a*_| represents its size. Type B and C scores were similarly defined as

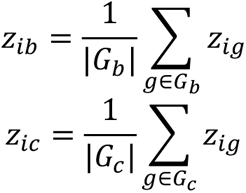

These mean z-scores were normalized to be bounded between [0, 1].

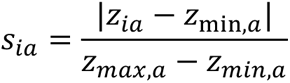

Where *z*_*min*,*a*_ is defined as the 40^th^ percentile of *z*_*ia*_ among L2/3 cells and *z*_*max*,*a*_ is defined as the 95^th^ percentile. Next, we normalized these scores such that they sum to 1,

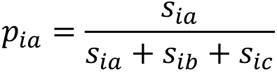

These scores can be directly visualized at single-cell level using an additive blending between three (archetypal) colors. To assign discrete cell type labels, one can also assign a cell to the type that has the highest ABC scores. Cells with clear A, B or C identity, i.e. archetypal cells, are defined as those whose highest ABC scores were greater than 0.6 (max(*p*_*a*_, *p*_*b*_, *p*_*c*_) > 0.6).

### Pseudotime analysis for the L2/3 transcriptomic continuum

We computed pseudo-temporal coordinates for L2/3 cells to understand their continuous organization. We first computed the top 50 PCs using z-scored L2/3 type-identity genes, and generated a k-nearest-neighbor (k=50) graph between cells in the PC space. The graph was built using the function *Scanpy.pp.neighbors*. Next, we ran a diffusion map (19) using *Scanpy.tl.diffmap* and ran diffusion pseudotime using *scanpy.dpt* following default parameters. The cell with the smallest PC1 was assigned as the as root cell and serves as the starting point of diffusion. As a result, each cell was assigned a pseudo-temporal coordinate, and cells were ranked by the increasing order of pseudotime.

We also ranked L2/3 type-identity genes according to their expression along the pseudotime coordinates. We defined the pseudotime of a gene (*T_g_*) as the weighted average of cell pseudotime,

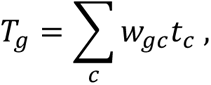

where *t*_*c*_ is the pseudotime of the cell (*c*), and *w*_g*c*_ is the weight of the cell *c* contributing to the gene *g*. The weight of a gene is defined by its expression level and sums to 1,

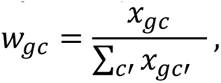

where *x*_g*c*_ is the size and log normalized expression. This gene pseudotime was defined at P28NR and was kept fixed to give a consistent representation for comparisons between conditions.

### Archetypal analysis (the multi-tasking framework)

We used the python package *py_pcha*: https://github.com/ulfaslak/py_pcha to determine the geometry of the L2/3 cell type continuum in the PC1-PC2 space derived from the L2/3 type-identity genes. The tool uses archetypal analysis, also known as the principal convex hull analysis, to infer a triangular boundary to the L2/3 transcriptomes. The same procedure (with parameters *delta=0 and noc=3*) was applied to different combinations of data (snRNA-seq, MERFISH, different conditions, harmonized data, and shuffled data; see main text).

We evaluated the significance of the triangular fit by t-ratio test as proposed by the multi-tasking framework (10). T-ratio is the ratio between the area of the convex hull of the data and that of the principal convex hull (the triangular bound in our case). We calculated the area of the convex hull using the python package *scipy* and its function *scipy.spatial.ConvexHull*. We tested the significance of the observed t-ratio by comparing it to t-ratios of shuffled data by permuting each gene independently across all cells. P-value was computed based on 1000 shuffles.

### Simulations of continuous and discrete types

First, we describe our procedure to simulate transcriptomic continua. Let cells and genes be ranked, such cells with a particular ranking express genes with a similar ranking. Let *i* be an integer representing the cell ranking, which ranges from 1 to *N*_*c*_, with *N*_*c*_ denoting the total number of cells. Let *j* be an integer representing the gene ranking, which ranges from 1 to *N*_g_, with *N*_g_ denoting the total number of genes. We use 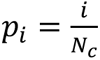 and 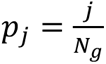 to represent the normalized rankings such that both the gene rankings and cell rankings range from 0 to 1. We consider a model where the expected expression level of gene *j* in cell *i* is a Gaussian such that:

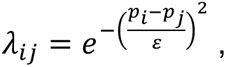

where *ε* denotes the level of noise. Gene counts are then drawn from a scaled Poisson distribution parametrized by *λ*_*i*j_:

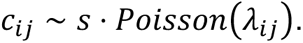

For our simulation, we chose *N*_g_ = 60, *N*_*c*_ = 600, *s* = 100, with different *ε* values between 0.1 and 1. Random numbers were generated using the python package *numpy.random*.

Second, for simulating discrete types, we assumed a model wherein each type is distinguished by a set of “marker” genes. Let *i* be a cell and *C*(*i*) be the cell type it belongs to. Let *j* be a gene and *C*(*j*) be the cell type of which it is a marker. The expected expression level of gene *j* in cell *i* was modeled by a binary matrix:

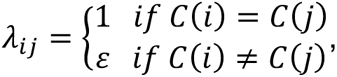

where *ε* denotes the amount of leaky expression, i.e., the level of noise. *ε* takes values between 0 and 1. The larger *ε* is, the noisier the types are. The actual count matrix was drawn from a scaled Poisson distribution parametrized by *λ*_*i*j_,

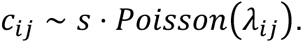

We simulated three discrete types (A, B, C), each with 20 marker genes and containing 200 cells, with different *ε* values between 0.1 and 1. To assign an order between types, among the 20 marker genes, 6 were shared between neighboring types, such that 6 type-A markers were also expressed in type B, and 6 type-B markers were also expressed in type C, and so on.

### Aligning V1 coordinates across sections

In our samples, V1 spanned about 2 mm (or ∼2,000 microns) along the tangential (M-L) axis of the cortex, with different sections having different lengths as defined by the expression domains of V1 marker genes *Scnn1a* and *Igfbp4* (see Main text). To quantify the *in situ* patterns of specific genes within V1 and in other cortical regions across samples, we defined a normalized tangential coordinate system relative to the length of V1. Specifically, for each section we used the tangential locations of the two ends of V1, *t*_*m*_ and *t*_*l*_, to normalize tangential coordinates along the cortex:

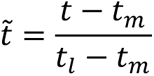

In this normalization, 0 and 1 represent the two ends of V1 (medial-most and lateral-most) respectively. And values <0 and 0>1 represent regions outside of (flanking) V1. Expression levels of specific genes are quantified according to this normalized coordinate system.

### Focused clustering

In addition to the cell types identified previously, we applied K-means clustering (*sklearn.cluster.Kmeans)* using only L2/3 type-identity genes as features, to identify focused clusters. As mentioned in the main text, the motivation of this analysis is to uncover L2/3 cell types in DR while ignoring changes to other gene programs related to cell states. This procedure generates matched cell types between NR and DR.

### Principal component analysis (PCA)

We applied PCA using *sklearn.decomposition.PCA.* In **Figure 3**, we applied it on L2/3 glutamatergic neurons combined from both P28NR and P28DR mice, using a set of unbiasedly identified highly variable genes (HVGs) as features. For HVGs identification, we considered n=21,222 genes that had non-zero counts in at least 10 cells. For each gene, we computed the variance-mean ratio on counts-per-10k-transcripts (CP10k) normalized data. Under a Poisson distribution, this variance-mean ratio is expected to be a constant. Indeed, this value is stable across orders of magnitude difference in mean expression across genes. To select HVGs with different baseline (mean) expressions, we grouped genes into decile-bins according to mean expression, and for each bin we selected the top 30% genes with the most variance-mean ratio. N=6,360 genes were selected, and most (n=270/286) L2/3 type-specific genes are part of the HVGs.

In **Figure 4**, we applied PCA using L2/3 type-identity genes (n=286; **Table S1**) as features. Focusing on L2/3 type-identity genes, rather than HVGs, allowed us to study the impact of visual deprivation on the L2/3 type-identity gene programs, while ignoring changes to other gene programs.

We evaluated the degree of overlap between two PCA eigenvectors *v*_*i*_ and *v*_j_as the absolute value of their dot product (S*v*_*i*_ ⋅ *v*_j_S). As the eigenvectors are orthonormal, this value is bounded between 0 and 1.

### Gene ontology (GO) analysis

We used the tool *EnrichR*: https://maayanlab.cloud/Enrichr/ for GO analysis. All enrichment analyses were performed by comparing a gene list of interest with a background set of top 5000 most expressed genes in L2/3 (P28NR). Top 10 enriched GO terms of Biological Process (2023 version) that were statistically significant with FDR < 0.05 were shown. For terms belonging to the same branch (parents or children), only one term with the highest FDR was shown in the main figures to reduce redundancy. Terms annotated as “obsolete” according to quickGO (https://www.ebi.ac.uk/QuickGO/annotations) were also removed from the main figures. All enriched terms (unfiltered) were shown in supplementary figures.

### Previously identified L2/3 stimulus-response genes

Hrvatin et al. (11) reported 611 stimulus-response genes identified from different cell types in V1, among which we filtered out those that are related to L2/3 glutamatergic neurons (labeled as “ExcL23” in Ref (48)), including 42 early-response genes and 37 late-response genes. These genes are listed in **Table S5**.

We used the Fisher exact test with n=6,360 HVGs as the background to test for enrichment or depletion of the overlap between PC-driving genes and the above activity-regulated genes.

### Quantification of cell redistribution in DR

We profiled the cell-density distributions of L2/3 cells in PC1-PC2 space derived from type-identity genes using the python package *seaborn.histplot*. The distributions were robust across a range of bin widths, and we used bin-widths of 1∼1.5 for visualization. We used the Jensen-Shannon (JS) divergence (49) to measure the level of difference in cell-density distributions between NR and D and across samples. This metric was calculated using *scipy.spatial.distance.jensenshannon*.

Optimal transport analysis was performed using the python package *POT*: https://pythonot.github.io/. The program computes the optimal transport map (28, 50) connecting the empirical distributions of NR cells and DR cells in PC1-PC2 space derived from type-identity genes. We first used *ot.dist* to compute pairwise distance matrix between NR and DR cells with default parameters. We then used *ot.emd* to calculate the transport map. The result was a matrix whose elements denote transport couplings between each pair of NR and DR cells, a proxy for their transcriptomic correspondence. We visualized this result by a coarse-grained vector field as follows. First, the PC1-PC2 space was meshed into equal-sized 2D bins. For each 2D bin, we computed a vector representing the mean moving direction and magnitude for all cells that fall into the bin. The vectors are visualized by arrows whose direction represents the average moving directions among local cells, whose lengths are proportional to the movement magnitudes, and whose color (darkness) is proportional to the number of cells represented by the bins.

### Identification of differentially expressed genes (DEGs) between NR and DR

In scRNA-seq analysis, it is common to treat individual cells as samples in statistical tests for identifying DEGs. This approach often leads to many statistically significant genes as even modest effect sizes can appear unlikely (low p-values) under the “no effect” null hypothesis owing to large cell numbers. Therefore, as a conservative measure to identify DEGs between NR and DR, we chose to regard the biological samples, rather than single cells, as independent data points.

We compared NR vs DR across independent biological samples, with each condition having four independent biological samples: P28 rep1, P28 rep2, P38 rep1 and P38 rep2. Raw counts from cells of the same types and samples were aggregated to produce pseudo-bulk profiles. We only considered genes with mean expression (in counts per million transcripts; CPM) > 10 in at least one type in either NR or DR. We performed independent sample t-test (*scipy.stats.ttest_ind*) on the pseudo-bulk profiles between NR and DR for each type. The resulting p-values were adjusted by the Benjamini-Hochberg procedure to calculate the false discovery rate (FDR). Effect sizes were quantified as the log2 fold change (in CPM) in DR compared with NR. DEGs were defined as those with *FDR* < 0.05 and |log_2_ *FC*| > 1.

## Data and code availability

- MERFISH data generated by this study was deposited on Zenodo with DOI: <u>10.5281/zenodo.13916878</u>
- snRNA-seq data was downloaded from GEO with the accession number <u>GSE190940</u>.
- Code to reproduce L2/3 type specific genes (Table S1, n=286) was obtained from the GitHub repository: https://github.com/shekharlab/mouseVC
- Code to reproduce the figures in the manuscript was deposited in the following GitHub repository: https://github.com/FangmingXie/vision_and_visctx

## Acknowledgments

The authors thank members of the Zipursky lab for critical feedback. This work was supported by a grant from the W.M. Keck Foundation to S.L.Z, a NIH grant 1F31 NS131016 to S.B., the Hellman Foundation (K.S.) and McKnight Foundation (K.S.). K.S. is an investigator with the Glaucoma Research Foundation and is supported by the Melza M. and Frank Theodore Barr Foundation. S.L.Z. is an investigator of the Howard Hughes Medical Institute.

## Author contributions

F.X., S.J., R.X., K.S., and S.L.Z. conceived the project. R.X., Z.T. and X.X. generated the MERFISH data. F.X. led the analyses. S.J., S.B. and K.S. contributed to analyses. F.X., S.J., S. B., K.S. and S.L.Z wrote the paper.

